# Modeling myosin Va liposome transport through actin filament networks reveals a percolation threshold that modulates transport properties

**DOI:** 10.1101/2021.08.10.455842

**Authors:** S. Walcott, D. M. Warshaw

## Abstract

Myosin Va (myoVa) motors transport membrane-bound cargo through three-dimensional, intracellular actin filament networks. We developed a coarse-grained, *in silico* model to predict how actin filament density (3-800 filaments) within a randomly oriented actin network affects fluid-like liposome (350nm vs. 1,750nm) transport by myoVa motors. 5,000 simulated liposomes transported within each network adopted one of three states: transport, tug of war, or diffusion. Diffusion due to liposome detachment from actin rarely occurred given at least 10 motors on the liposome surface. However, with increased actin density, liposomes transitioned from primarily directed transport on single actin filaments to an apparent random walk, resulting from a mixture of transport and tug of wars as the probability of encountering additional actin filaments increased. This phase transition arises from a percolation phase transition at a critical number of accessible actin filaments, *N*_*c*_. *N*_*c*_, is a geometric property of the actin network that depends only on the position and polarity of the actin filaments and the liposome diameter, as evidenced by a five-fold increase in liposome diameter resulting in a five-fold decrease in *N*_*c*_. Thus, in cells, actin network density and cargo size may be regulated to match cargo delivery to the cell’s physiological demands.

## Introduction

In eukaryotic cells, intracellular cargo transport, such as insulin granules that are destined for secretion at the cell membrane, is shared by molecular motors that travel along the microtubule and actin cytoskeletal highways [1]. Beginning at the Golgi where secretory vesicles are formed, kinesin motors provide longrange transport along microtubules oriented towards the cell membrane. Final short-range transport and delivery to the cell membrane relies on vesicle hand-off to myosin Va (myoVa) motors that maneuver their cargo through the dense, randomly oriented cortical actin filament network [2–5]. However, the precise role of myoVa in cells is still debated: whether it’s a short-or long-range transporter for cargo delivery to a targeted location or whether myoVa tethers cargo to the actin network in close proximity to the cell membrane [5–9]. Are these putative myoVa roles compatible with an apparently random network of actin filaments with which the motors are engaged or does the structure of the actin filament network itself dictate which functional role myoVa motors adopt? The importance of these questions is underscored by genetic mutations in myoVa that lead to mis-localized cargo in cells [10] and neurological defects in humans [11, 12].

To define the challenges that myoVa motors face when navigating cargo through the cell’s three-dimensional (3D) actin filament network, investigators have built complexity *in vitro* within model transport systems, ranging from single myoVa transport, as the motor steps towards the plus-end of a single actin filament, to more complex liposome transport by teams of myoVa motors through 3D actin filament networks [13–20]. Understanding the molecular bases for transport outcomes in these *in vitro* experiments has relied on mechanistic mathematical modeling. These models incorporate numerous aspects of myoVa molecular structure and function but most importantly the force-dependent slowing of myoVa’s stepping rate when the motor experiences a hindering force applied externally to the cargo or due to a “tug of war” between motors on the same cargo that interact simultaneously with separate actin filaments [18,21,22]. *In silico* modeling of this latter scenario was critical to our previously interpreting the trajectories of liposomes transported by a myoVa team through 3D actin networks [19,20], providing insight into how the density and orientation of the local actin network surrounding the liposome dictated the motors’ short-range (∼ 1 *µ*m, 1-10 seconds) modes of transport motion (i.e. directed, diffusive-like, or stationary). However, these previous *in vitro* studies were limited experimentally, since exploring the vast experimental parameter space, such as the effects of cargo diameter and actin network density, was both technically and analytically demanding. Thus, there remains a critical gap between these *in vitro* experiments and myoVa’s function in cells, where cargo transport to a destination can occur over longer spatial and temporal scales (∼ 10s of *µ*m, minutes).

Mathematical models can help bridge the gap between the molecular and larger intracellular scales. Models keep track of a multitude of molecular-scale processes that occur simultaneously and impact each other, often leading to emergent properties (e.g. phase transitions). For example, phase transitions in the macroscopic mechanical properties of cross-linked actin networks in the presence of myosin-II filaments, i.e. whether the system flows like a fluid [23–25], remains solid like a gel [26, 27] or contracts into one [28] or more [29, 30] dense clusters, can be understood through mathematical modeling [31–33]. Here we use *in silico* modeling to investigate how myoVa-based cargo transport through a random 3D actin filament network over spatial and temporal scales observed in cells (∼ 10s of *µ*m, minutes) is affected by actin filament density within the network and by cargo diameter. We find that such long-range myoVa-based transport shows a cargo size-dependent phase transition as actin density is varied. Specifically, at low actin density, transport is rapid and unidirectional. In contrast, above a critical actin density that scales inversely with cargo diameter, transport is interrupted by pauses and loss of directionality, as motor teams undergo tug of wars and switch filaments, respectively. Cells may take advantage of this phase transition to switch roles between myoVa acting as a long range transporter or tether.

## Results

### Model of Long-Range Liposome Transport

In this study we used *in silico* modeling to define how long-range vesicular transport by myoVa motors through an actin filament network would be impacted by changes in actin filament density and cargo diameter. The foundation for this work was our previously developed and experimentally-validated, *in silico* mechanistic model for the transport of 350nm (radius *r*_*L*_ = 175nm), fluid membrane liposomes by teams of 10 myoVa motors through actin filament networks [19, 20]. However, this model probed short-range myoVa transport (∼ 1 *µ*m, 1-10 seconds) within a limited actin filament network (i.e. several actin filaments) rather than transport over spatial and temporal scales observed in cells (∼ 10s of *µ*m, minutes). Therefore, using the same liposome-motor complex, we modeled transport through a series of complex actin filament networks confined within a 20*µ*m diameter sphere (Fig. 1). These actin networks consisted of randomly placed filaments of varying number; covering roughly three orders of magnitude (i.e. 3 -800 filaments, see Supplementary Movies S1-S6). Since all networks were confined to the same diameter sphere, the number of actin filaments within the sphere can be used interchangeably with actin density. For each actin filament network, we simulated 5,000 liposome trajectories that all started with the liposome bound to an actin filament at the network center (Fig. 1A, t=0s). A trajectory concluded when either the liposome reached the boundary of the sphere or 100 seconds elapsed. In total, we simulated 1,040,000 liposome trajectories for the various model conditions. However, given our experimental parameter space, using our previous model algorithms was too computationally expensive. Therefore, we implemented a coarse-grained model, described below, that allowed us to more efficiently simulate myoVa-based liposome transport over larger spatial and temporal scales.

**Figure 1:**
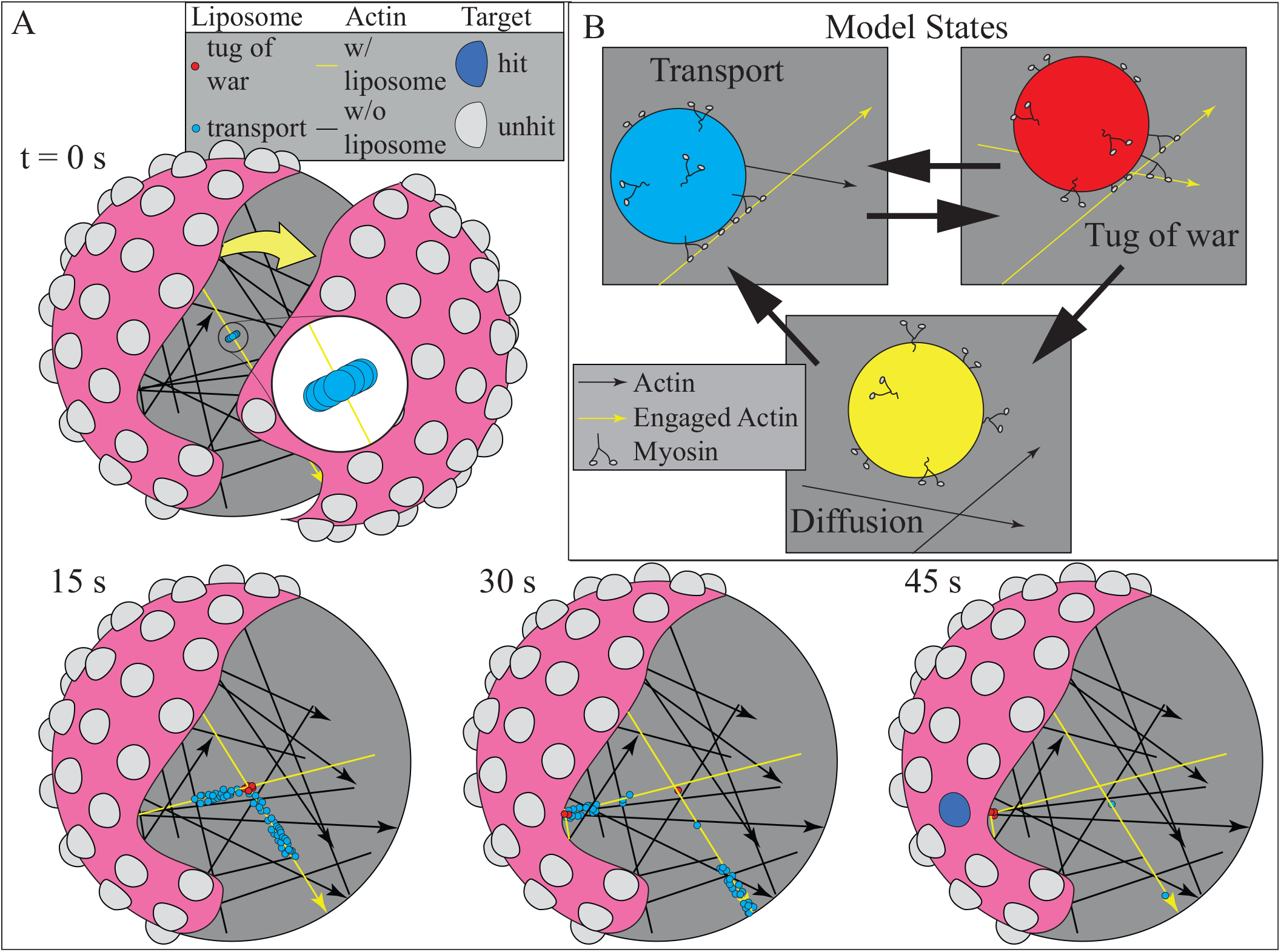
Schematic of the simulations. A. A random mesh of actin filaments span a spherical region (partially cut away for visualization purposes) of radius 10*µ*m (magenta). Fluid liposomes, of 350nm diameter, are transported (blue) by 10 myoVa motors, undergo tug of wars (red), or diffuse according to a coarse-grained model. 2*µ*m diameter spherical targets (gray) are distributed evenly over the surface of the sphere, and are hit (blue) if touched by a liposome. A simulation of 100 liposomes navigating a network of *N* = 25 actin filaments is shown (see Supplementary Movie S7 for a movie of the simulation). Actin filament thickness is exaggerated for visualization purposes; arrows indicate plus ends. B. Schematic of state transitions in the model. Liposomes in the transport state (blue) have myoVa motors engaged with a single actin filament. If a motor engages with a second actin filament, a tug of war (red) is initiated. Tug of wars may be resolved with motors remaining engaged with the original filament (straight outcome), switching to the new filament (turn outcome), or detaching. In the latter case, the liposome enters the diffusion state (yellow). The coarse-grained model used in our simulations does not keep track of the myoVa motors, but is validated by simulations of a more detailed model that does.

In this coarse-grained model (see Methods and Supplemental Material (SM) for details), the liposome is in one of three states at any point in time (Fig. 1B), undergoing: 1) transport, where the liposome-motor complex is engaged with and traveling along a single actin filament; 2) tug of war, where motors on the same liposome interact simultaneously with separate actin filaments or; 3) diffusion, where the liposome freely diffuses with an assumed diffusion constant for a 350nm sphere in water at 25^*°*^C (*D* = 1.3*µ*m^2^*/*s). A liposome in the transport state travels towards the actin filament plus-end along a spiral path dictated by the actin filament helix [34]. Once the liposome encounters a second actin filament within reach of the motors on its surface, it may transition into the tug of war state. When in a tug of war, the liposome is stationary until the tug of war is resolved when either: 1) motors engaged with one actin filament overcome the force generated by motors engaged with the other actin filament, thus returning the liposome to the transport state on the filament associated with the winning team of motors, or; 2) all motors detach from their engaged actin filaments and the liposome transitions to the diffusion state. When in the diffusion state, the liposome can only transition to the transport state after it collides with an actin filament. Rate constants between states depend on the proximity of actin filaments to the liposome, while the tug of war duration depends on the position and polarity of the actin filaments with which the motors are engaged. The rate constants for liposome state transitions were determined by fitting this coarse-grained model to thousands of simulations of our previous, more detailed model [19, 20] (see SM for the fits). Once the coarse-grained model’s predictive capacity was verified against the short-range predictive capacity of our previous model, long-range liposome trajectories in the various actin network models were simulated with a Monte-Carlo method (Fig. 1A; see SM for more algorithm details). Steric interactions between the liposome and the actin network are not included in this model, so we ensured that all actin networks had pore sizes larger than the liposome diameter (see SM).

### Liposome transport simulations at varying actin network density

Simulated liposome trajectories at low actin density (3 -50 filaments; Movies S1-S4) demonstrate that most liposomes move along individual actin filaments in the transport state (blue liposome in movies and Fig. 1; Fig. 2B) and thus rarely engage in a tug of war (red liposome in movies and Fig. 1B). By contrast, at high actin density (100 -800 filaments; Movies S5 and S6) most every liposome was within reach of multiple actin filaments and thus switched between transport and tug of war states (red liposome in Fig. 2C); resulting in random walk-like trajectories, as liposomes often switched filaments once the tug of war was resolved (Fig. 2C). This apparent change from primarily long-range, directed transport to more circuitous trajectories with increasing actin filament density suggests that network density triggers a shift in the liposome state distribution. Plotting the coarse-grained model state distributions as function of actin density confirms (Fig. 3) that the transport state predominates at low actin density, transitioning to a mixture of transport and tug of war states with actin densities greater than 50 filaments. Due to multiple motors on the liposome surface being simultaneously engaged with one or more actin filaments, liposomes rarely detached from actin to enter the diffusion state (yellow liposomes in movies and Fig 1; Fig. 2A) at any actin density.

**Figure 2:**
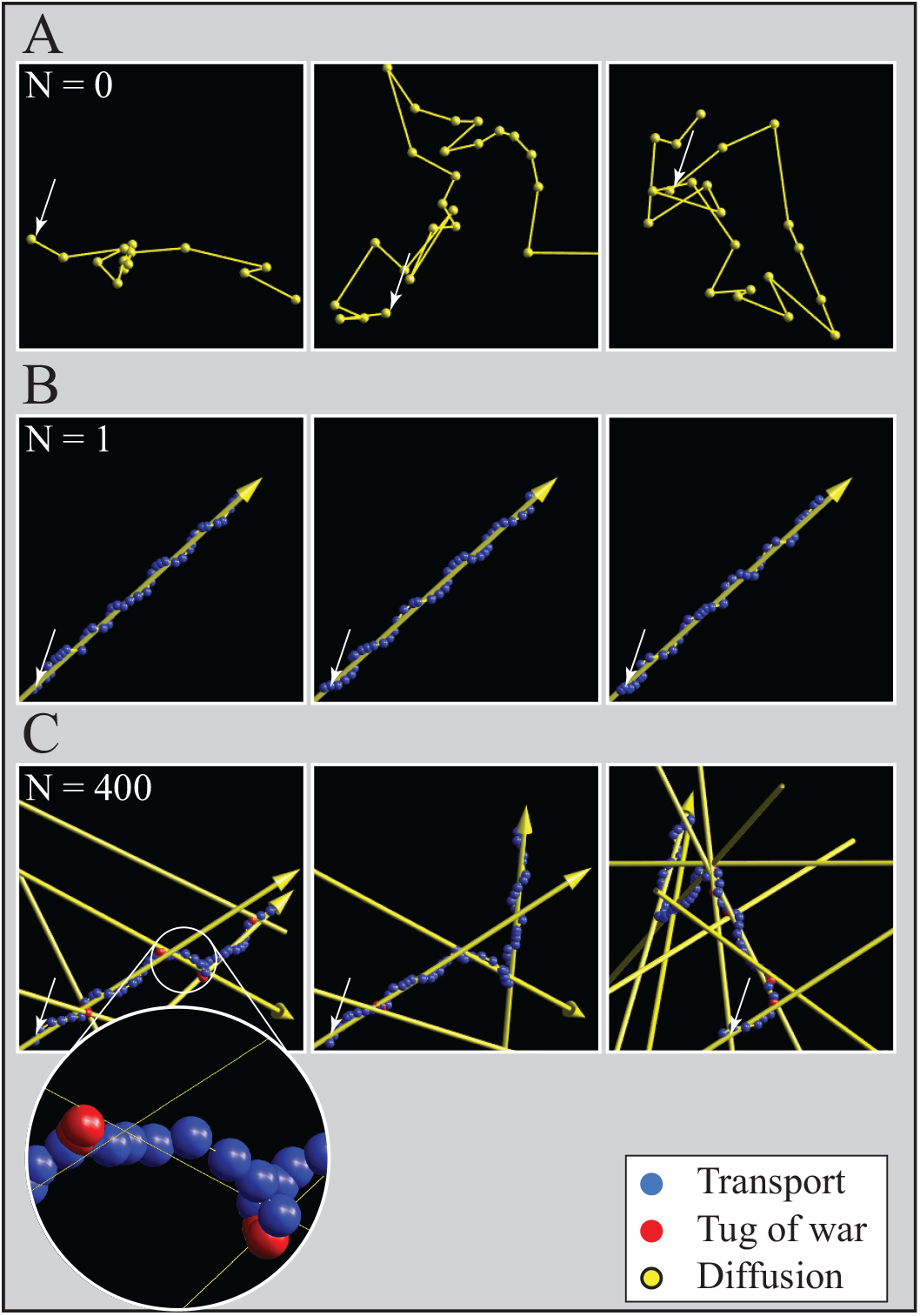
Increasing actin network density causes a change in liposome motion. A. Liposomes in the diffusion state (yellow spheres) describe a random walk. This is rarely observed in our simulations at any actin density. B. Liposomes transported along a single actin filament (yellow arrow with arrow head indicating plus-end) adopt the transport state (blue spheres). This is observed in our simulations at low actin density. C. At high actin density, liposomes transition between transport (blue spheres) and tug of war (red spheres) states, resulting in a random walk as they switch actin filaments once the tug of war is resolved. Actin filament thickness is exaggerated for visualization purposes (the magnification call-out in C shows actual size). Liposome position is shown at equally spaced intervals of Δ*t* = 0.75s, and Gaussian random noise of standard deviation *σ* = 10nm is added for visualization purposes. Small white arrows indicate liposome starting position.

**Figure 3:**
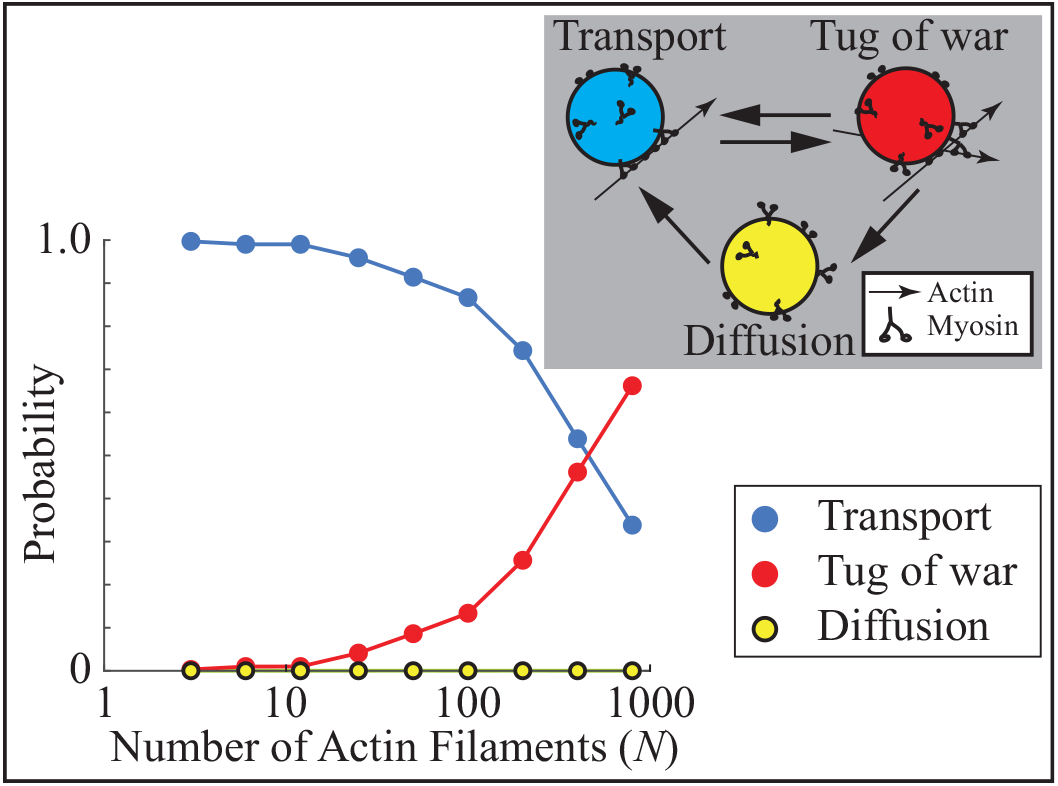
Actin network density triggers a shift in liposome state distribution. The average state of a simulated liposome shifts from being almost exclusively transport at low actin density to a mix of transport and tug of war at higher density. There is very little diffusion at any density. Inset shows a diagram of the model states and transitions. Error bars (SEM) are smaller than the symbols.

To relate how liposome state distributions dictate simulated transport characteristics, motion parameters typically used to describe particle trajectories were determined at each actin density. The average liposome speed (linear distance between initial and final position divided by simulation time; Fig. 4A) remains constant and equal to that for transport along a single actin filament (350nm/sec) at low actin density, but then begins to slow at actin densities *>* 50 filaments to a minimum of 53nm/sec at the highest density (i.e. 800 filaments). This slowing is explained by liposomes transitioning to a mixture of transport and tug of war states at high actin densities. The increased frequency of stationary periods during the tug of wars and the less-directed trajectories due to filament switching both contribute to the slower average speeds (Figs. 2C, 4A).

**Figure 4:**
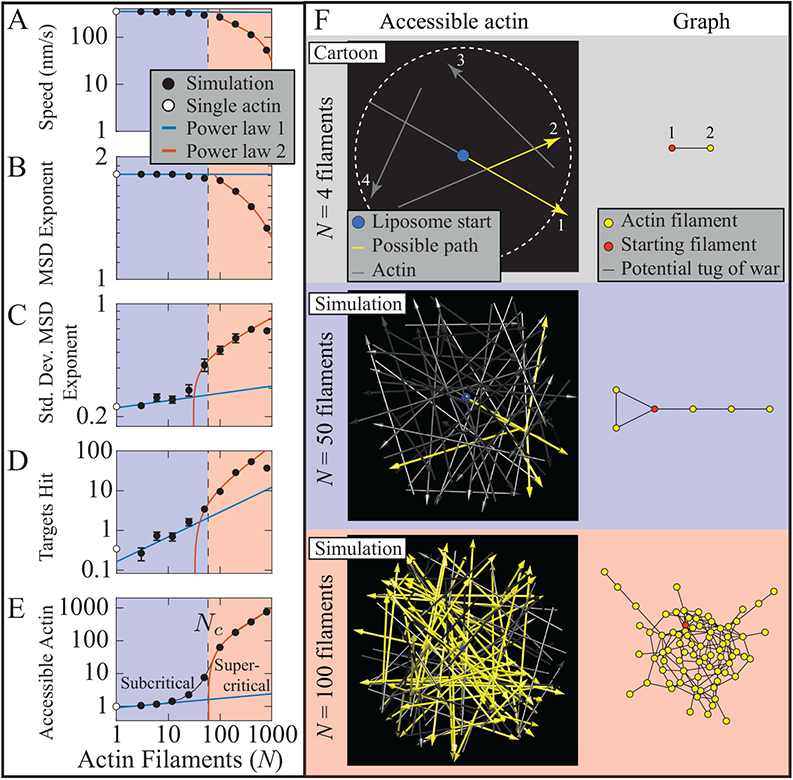
A change in the number of actin filaments accessible for transport underlies a phase transition in myoVabased transport with increasing actin density. Different quantities for liposome motion, i.e. liposome speed (A), mean squared displacement exponent (B; MSD), Std. Dev. of MSD (C), and number of targets hit by liposomes at the network boundary (D) show a change in scaling as best described by two different power laws similarly to the change in scaling that occurs in the number of accessible actin, *N*_*c*_ ≈ 60 actin filaments (E) (see text for details). The blue shaded region is where power law 1 applies, and the red shaded region is where power law 2 applies for the accessible actin. Error bars show SEM. White circles show values for a single actin filament. Schematics of accessible actin (F), represented as the number of potential transport paths on a given actin network (left), and as a graph (right). Top shows a cartoon with 4 actin filaments as a demonstration. In this network, actin filaments 1 and 2 are within reach of a myoVa on the liposome surface, *r*_*B*_ (i.e. distance from liposome center to the tip of myoVa on the liposome surface; see SM for details), at their nearest point, while actin filaments 1 and 3, and 2 and 3 are further than *r*_*B*_. Actin filament 4 is not accessible even being closer than *r*_*B*_ to filament 1 due to the liposome starting at the center of filament 1 and then travels towards the plus-end away from filament 4. A much greater percentage of the actin filaments are accessible for the larger network (*N* = 100 filaments, 78*/*100 = 78%) than for the smaller network (*N* = 50 filaments, 6*/*50 = 12%). In each graph (right), accessible actin filaments are shown as vertices (yellow circles, starting filament as red circle), and potential tug of wars as edges (black lines). Actin filament thickness is exaggerated for visualization purposes; arrow heads indicate plus-ends; see Supplementary Movies S8 and S9 for simulations of liposomes transported through the two networks.

Another transport descriptor is the mode of trajectory motion, defined by the scaling exponent of the mean squared displacement as a function of time (*α*_*MSD*_), where: *α*_*MSD*_ = 0 for a stationary liposome; *α*_*MSD*_ ∼ 1 for diffusive-like motion, and; *α*_*MSD*_ = 1.8 for purely directed, straight line motion, based on simulations (see SM). Therefore, at each actin density we calculated *α*_*MSD*_ for the entire simulated trajectory of each liposome (Fig. 4B; see Methods and SM for details). Liposomes primarily adopt a directed mode on low actin density networks (i.e. *<* 50 filaments), shifting toward a more diffusive-like mode at higher densities to a minimum of *α*_*MSD*_ = 1.34 at the highest density (i.e. 800 filaments). This decrease in *α*_*MSD*_ is consistent with the transitions in the liposome state distributions described above, with stationary tug of wars and a loss of directionality due to filament switching contributing to the overall motion appearing more diffusive-like; however, calculating *α*_*MSD*_ over the entire trajectory averages together times when an individual liposome might adopt different modes of motion. Calculating *α*_*MSD*_ over a much shorter time scale (one second) shows that on the dense actin networks (i.e. *>* 50 filaments), liposomes adopt a mixture of directed and stationary modes of motion (see SM), which directly reflects the transitions in the transport state distributions described above. Thus, this shift in liposome motion with actin density is emphasized both in the average *α*_*MSD*_ for the entire trajectory or in the standard deviation of the *α*_*MSD*_ calculated over one second of the trajectory (Fig. 4C), where a small standard deviation indicates a single predominant mode of motion, while a large standard deviation indicates a mixture of modes.

### Metric for quantifying global liposome transport

Simulations described above identified a transition in transport characteristics as actin network density increased. However, these motion parameters are average characteristics that describe the transport of a single liposome. If myoVa intracellular transport is potentially associated with cargo delivery to targeted locations, then this shift in trajectory motion characteristics with actin filament density may be a regulated cellular property that can modulate targeted cargo delivery. Whether transport is considered targeted for a given network is best described as a property of a liposome population, rather than of a single liposome. We, therefore, defined a metric for targeted liposome delivery by evenly spacing 100 spherical (1*µ*m radius)”targets” on the network boundary surface (Fig. 1A). This spatial distribution translates into 35% of the total boundary surface being composed of target sites that can be “hit” by a delivered liposome. Thus, at sparse actin filament densities where liposomes may travel from the network center to the network boundary along a single actin filament, the probability of hitting a target is 0.35 on average. In contrast, at higher filament densities, where a multitude of potential paths from the network center to the surface are available, then, most if not all, of the 100 targets may be hit. Thus, the smaller the number of targets hit, the more directed the transport on a given network. This metric of liposome transport (i.e. targets hit) provides a simple quantifiable measure for how a liposome population would be delivered within a given actin filament network, so that changes to liposome delivery can be readily detected as a result of variations in actin filament density and liposome diameter.

### Liposome transport undergoes a phase transition at a critical accessible actin filament density

Model simulations demonstrate that as actin filament density increases, so too does the number of targets hit (Fig. 4D). This relationship is best described by two different power laws, which are apparent by plotting the data on a log-log scale. Specifically, at low density, the number of targets hit increases slowly, while increasing more rapidly at higher density (Fig. 4D), until deviating at the highest actin filament density (i.e. 800 filaments), where liposome motion is so slow (see above, Fig. 4A) that liposomes generally do not reach the network boundary during the 100s simulation. The presence of two scaling regimes suggests a phase transition. In fact, all properties of transport we measured show a change in scaling that occurs at a critical actin density, *N*_*c*_ ≈ 60 filaments, defined as the *x*-intercept of the fit to the second phase (Fig. 4A-D). This phase transition may be related to the number of actin filaments accessible to liposome transport.

The number of accessible actin filaments is a geometric property of a given actin network. To be considered accessible to a liposome traveling along an actin filament, a second actin filament must be in reach of motors on the liposome surface. After simulating all potential paths that a liposome could traverse from its start at the network center to the network boundary without detaching from actin, this set of filaments was defined as the number of accessible actin filaments for a given network (see SM for more details, Fig. 4F for examples). The number of accessible actin filaments as a function of total number of actin filaments within the network is also described by two different power laws (Fig. 4E). The transition between these two phases occurs at the same actin filament density, i.e. *N*_*c*_ ≈ 60 filaments, as for the other transport properties (Fig. 4E). Therefore, at actin densities below this critical number (*N < N*_*c*_), the number of accessible actin is small with few potential paths for liposome transport (Fig. 4E) so that liposomes are predominantly in the transport state (Fig. 3). Whereas above this value (*N > N*_*c*_), the number of accessible actin increases rapidly, resulting in numerous, complex transport paths, due to a mixture of transport and tug of war states (Fig. 3) that culminate in random walk-like trajectories (Fig. 2C). Therefore, the phase transition in the number of targets hit is a direct reflection of the network geometry, i.e. a phase transition in the number of accessible actin filaments and potential paths as the liposome weaves its way through the network. Here we show that this geometric phase transition is a percolation phase transition, described in graph theory and physics.

### A percolation phase transition in the actin network underlies the phase transition in liposome transport

Percolation phase transitions, originally defined in the context of water draining through a porous medium [35], occur in a broad range of discrete networks, including the internet [36], power grids [37], epidemics [38], liquid to gel phase transitions in polymers [39–42] and actin networks in the presence of cross-linking proteins [43]. Common to all of these systems, and potentially the liposome transport system described here, is that a phase transition (i.e. a change in power law scaling) arises from a sudden increase in overall network connectivity as connection probability increases. As a demonstration of this process, consider a graph representing a network with *N* vertices and a line that connects a pair of vertices termed an edge (Fig. 5A). As additional edges connect pairs of vertices chosen at random, the average edges per vertex for the network, *z*, increases linearly from 0. However, the size of the giant component, i.e. the largest number of vertices that form a connected cluster as a fraction of *N* (Fig. 5A), varies discontinuously with *z* (Fig. 5B). When *z* is small, the giant component makes up a negligible fraction of the total number of vertices (Fig. 5B). Such a network is said to be *subcritical*. Beyond a critical *z* value, the percolation threshold, the giant component makes up a non-negligible fraction of the vertices and rapidly increases until all vertices are connected. Such a network is said to be *supercritical*. As the size of the network *N* increases, this phase transition becomes sharper and, in the limit of an infinitely large *N*, shows a discontinuity in slope at the percolation threshold, *N*_*c*_ (Fig. 5B). Though networks can differ in their details, all percolation phase transitions arise from a sudden increase in network connectivity, measured in the above example as the size of the giant component, which underlies different power law scaling on subcritical and supercritical networks.

**Figure 5:**
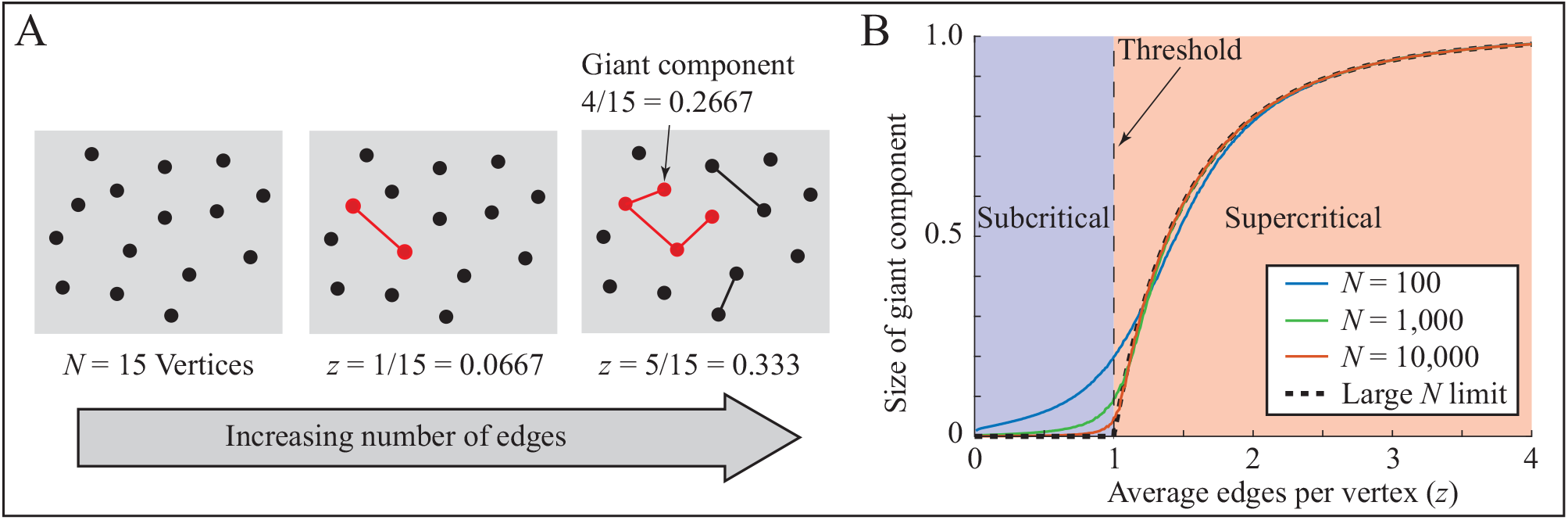
A schematic of a percolation phase transition on a network with random connections. A. A cartoon of *N* = 15 vertices (black circles), with an increasing number of edges, where an edge (line) connects two vertices. The giant component is the largest number of edge connected vertices (red circles and lines). *z* equals the average number of edges per vertex within the network. B. Plot of the size of the giant component as a proportion of the total number of vertices versus *z* for networks with different numbers of vertices, *N*. As *N* approaches infinity, the size of the giant component becomes non-zero above a critical number of edges, the percolation threshold, and networks switch from being subcritical to supercritical.

Assuming the transport characteristics of our modeled liposome transport system is reflective of a percolation phase transition, each actin filament can be represented as a vertex and an edge, as a tug of war, connecting two vertices (i.e. separate actin filaments) due to the filaments being spatially close enough so that myoVa motors on the same liposome can simultaneously interact with both filaments (Fig. 4F). Increasing the number of actin filaments in the network, *N*, adds vertices and, with each additional vertex the potential for an edge (i.e. tug of war) increases. If our system undergoes a percolation phase transition, then we expect to see a critical actin filament density, *N*_*c*_, above which a non-negligible fraction of the total vertices create a connected cluster and form the giant component. That is, above *N*_*c*_ a liposome starting on a filament at the network center could potentially reach a non-negligible fraction of the other actin filaments without having to detach and undergo diffusion. This quantity is what we have called the accessible actin. The fact that there is a change in scaling at *N*_*c*_ = 60 filaments for not only accessible actin but each transport property we measured (Fig. 4A-E) suggests that there is a phase transition in liposome transport within these networks that arises from a percolation phase transition in the actin network. However, the discontinuity at this threshold is not as sharp as might be expected (see Fig. 5B).

To better identify the percolation threshold, *N*_*c*_, by a clear discontinuity in the number of accessible actin filaments, we accounted for two factors that contributed to smoothing of this phase transition in Fig. 4E. The first of these factors is stochastic effects introduced by the random placement of filaments within each network. Therefore, we generated 10,000 different actin networks at each actin density below and above the *N*_*c*_ estimated from Fig. 4E. A histogram of the accessible actin as a proportion of the total actin on subcritical networks with *N < N*_*c*_ (e.g. *N* = 35, Fig. 6A) shows a single peak at the most common condition; a single accessible actin filament. This single accessible actin filament is the actin filament on which the liposomes start, and that trajectories along 2, 3 or more actin filaments are increasingly rare (Fig. 6A). However, as the network density crosses *N*_*c*_ (e.g. *N* = 70, Fig. 6B), the distribution of the accessible actin as a proportion of the total actin becomes bimodal, with multiple accessible actin filaments becoming more common, though a single accessible actin filament can still occur. Finally, at large supercritical network densities (i.e. *N > N*_*c*_, e.g. *N* = 125, Fig. 6C), the bimodal distribution is distinct, with a minor component at a single accessible actin filament still apparent. This bimodal distribution for supercritical network densities arises because the accessible actin must include the actin filament where transport starts. Due to the stochastic nature of each actin network, the starting actin filament may not be part of the network’s giant component. A single accessible actin filament is therefore possible, leading to the existence of a minor component of the distribution, i.e. mode 1. For each actin network density, *N*, the distributions of the accessible actin cluster size as a proportion of network density (Fig. 6A-C) were determined and these distributions fitted to define the mode(s) (see SM). By plotting these mode(s) of the fit versus *N* (Fig. 6D), an abrupt transition exists at the percolation threshold *N*_*c*_ = 58, where the actin network transitions from subcritical actin densities with predominantly single actin filament transport to supercritical actin densities where transport is on connected actin filament clusters (Fig. 6). This precise estimate of *N*_*c*_ supports our previous assumption that the apparent phase transition in the number of targets hit and accessible actin filaments that we estimated at *N*_*c*_ = 60 is a network property that reflects a percolation threshold and underlies transitions in all of the metrics used to describe liposome transport (Fig. 4A-E) as well as the liposome state distributions (Fig. 4D).

**Figure 6:**
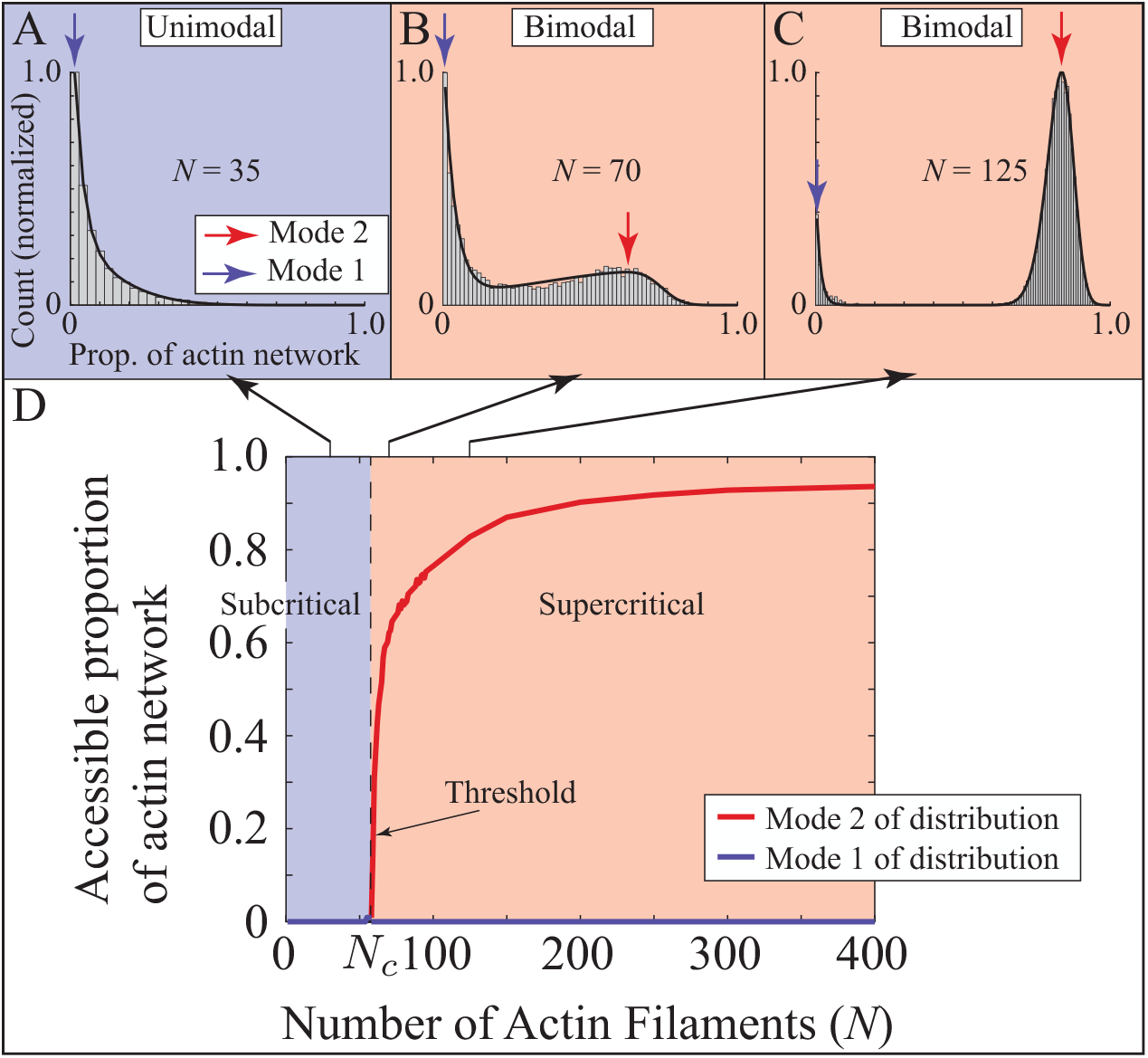
Percolation in accessible actin. A-C. Histograms of the proportion of accessible actin within 10,000 networks for various network densities. On a subcritical network (A, *N* = 35), the histogram has a mode of 1. On a supercritical network (B and C, *N* = 70 and 125, respectively), the histogram is bimodal. D. Plotting the mode(s) of the distribution as a function of the number of actin filaments shows the percolation threshold, *N*_*c*_ = 58, and the transition from subcritical to supercritical networks. Note that to calculate the accessible proportion of the actin network, we subtract 1 from accessible actin and divide by the total number of filaments.

### Percolation threshold is liposome diameter-dependent

The percolation threshold for liposome transport described here is a geometric property of the network system that involves an interplay between the actin network density and liposome diameter. Specifically, edges in the percolation network graph (Fig. 4F) occur only if motors on the liposome surface are within reach of nearby actin filaments so as to initiate a tug of war. Therefore, a dependence of *N*_*c*_ on liposome diameter should exist, which we tested by increasing the liposome diameter five-fold to 1,750 nm (*r*_*L*_ = 875nm) (see SM for details). Estimating the percolation threshold as before through histograms of the accessible actin cluster size as a proportion of a given network density (Fig. 6) resulted in an *N*_*c*_ = 11 filaments (Fig. 7A), a 5.3-fold decrease compared to that for liposomes of 350 nm diameter. This liposome diameter-dependent shift in *N*_*c*_ was similarly reflected in a ∼ 5-fold lower lower actin density at which phase transitions occurred for the coarse-grained model state distributions (Fig. 7B), and all other transport descriptors (Fig. 7C). These results confirm that the percolation threshold, *N*_*c*_, scales inversely with cargo diameter and underlies the phase transitions in liposome transport (see SM for further analysis), specifically from directed motion in subcritical networks to random walk-like motion interspersed with stationary pauses in supercritical networks.

**Figure 7:**
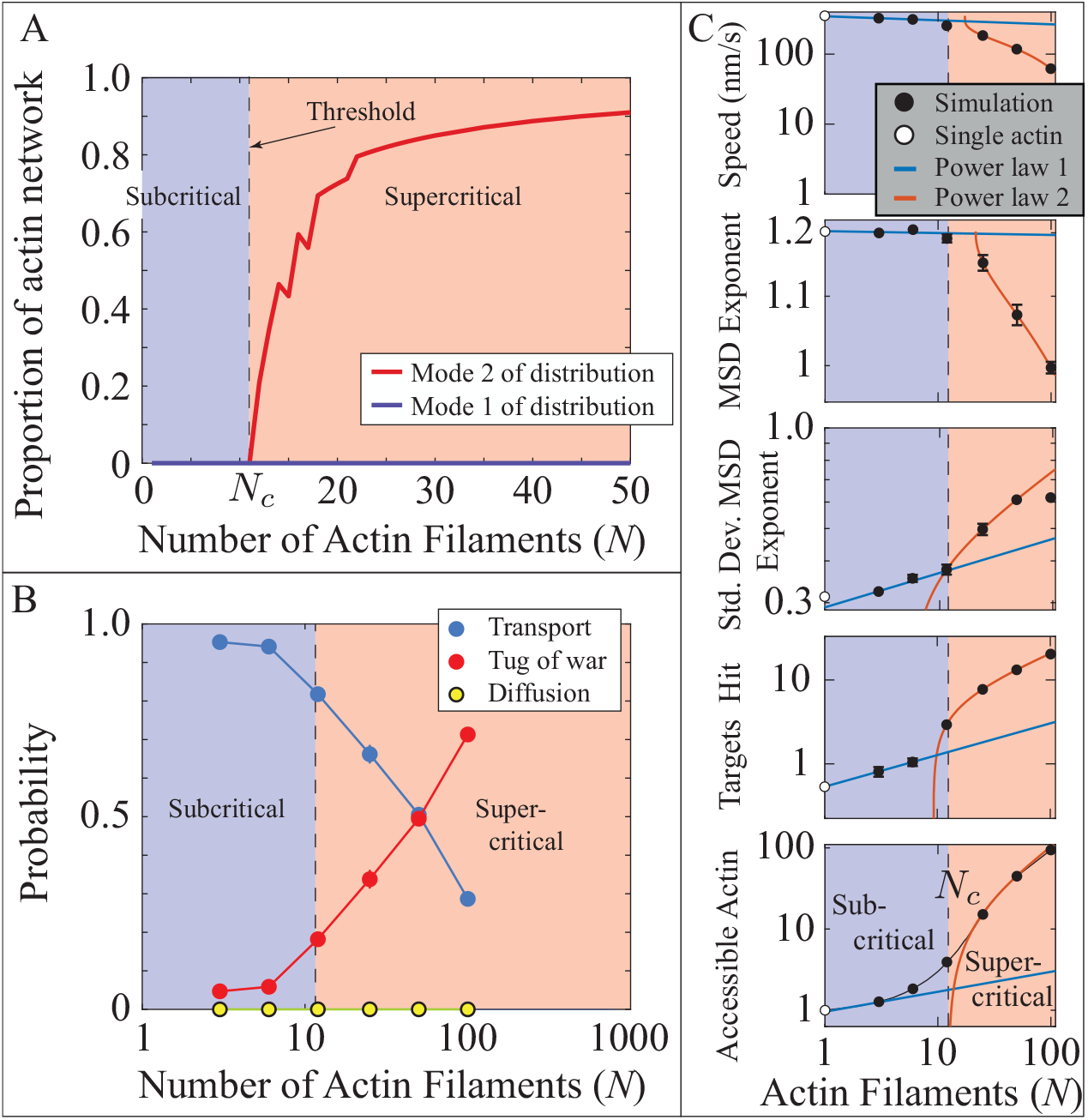
A five-fold increase in liposome size decreases the percolation threshold *N*_*c*_, and the transition between scaling regimes, by a factor of five. A. Histograms of accessible actin as a proportion of the total number of actin filaments within the network switch from being unimodal to bimodal at a critical network size of *N*_*c*_ = 11 filaments (see Fig. 7). B. The average state of a simulated liposome shifts from being almost exclusively transport to a mix of transport and tug of war as the number of actin filaments increases past *N*_*c*_. Error bars (SEM) are smaller than the symbols. C. Different quantities describing liposome motion (described in detail in main text) show a change in scaling similar to that which occurs in the number of accessible actin. In the plot of accessible actin (bottom graph). White circles show predicted values for a single actin filament. Each transition occurs at a network size roughly five-fold less than for smaller liposomes (see Fig. 7). Error bars show SEM.

### Liposome diffusion and its impact on transport outcomes

In simulations described above, we observed liposome trajectories in supercritical actin networks with mean squared displacement exponents approaching 1 (Figs. 4B and 7C). This diffusive-like mode of motion arises from liposomes being transported along a sequence of randomly-oriented actin filaments (Figs. 4 and 7B). In simulations at all actin densities, true diffusion is rare because there are sufficient motors on the liposome to engage one or more actin filaments so that liposome detachment does not occur. In a cell, however, diffusion may be initiated by deactivation of motors, the presence of actin-binding proteins that cause myoVa motors to detach, or myoVa-bound cargoes being assembled far from the actin network so that myoVa motors cannot engage. Therefore, we performed simulations as described above, but with liposomes starting at the network center unbound to actin (see Movies S10-S14).

As expected, on the lowest actin density networks (*N <* 50 filaments), liposomes were predominately in the diffusion state (Fig. S6B, S7D SM). However, on networks with *N >* 50 filaments, the model state distribution becomes nearly identical to simulations where liposomes start bound to actin (see SM). This transition from the diffusion state to a mixture of transport and tug of war states corresponds to the percolation threshold (*N*_*c*_ = 58). This, however, is a coincidence, since this transition does not correspond to the percolation threshold for the larger liposomes (*r*_*L*_ = 875 nm; Fig. S7E SM).

## Discussion

MyoVa transport of vesicular cargo along the cell’s actin cytoskeletal network involves a complex interplay between motors on the cargo surface that power cargo movement, the cargo dimensions, and the spatial arrangement of the actin filament tracks on which transport occurs. These three components (i.e. motors. cargoes, tracks) can vary significantly among cell types and even within a cell, as physiological demands warrant. With all three components potentially regulated, the cell then possesses a highly tunable cargo transport and delivery system. However, defining how each transport component contributes to cargo delivery requires investigators explore an enormous parameter space within cells that is experimentally challenging. To avoid these complexities, we developed a coarse-grained *in silico* model of liposome transport by myoVa motors within complex networks of randomly arranged actin filaments. Liposome movement transitions between three states of motion described as transport, tug of war, or diffusion, with the distribution of these states varying with actin density. Specifically, as actin density increases, liposomes transition from the highly directed “transport” state along single actin filaments to one described as a random walk, as the probability of a liposome encountering another filament increases and the motors on the liposome surface engage in a tug of war, with the resultant outcome a potential change in travel direction. This transition in transport characteristics arises from a percolation phase transition in the number of accessible actin filaments (Figs. 4E, 6D), which is a geometric property of the actin network that depends only on the position and polarity of the actin filaments and the liposome diameter. Interestingly, this percolation phase transition is predictive of phase transitions in various metrics of liposome transport that occur at the same actin density as the percolation threshold.

### Percolation transition in liposome transport through actin networks

Percolation transitions have been described previously in cytoskeletal networks in which the network consisted of actin filaments, actin filament cross-linking proteins, and myosin-II motor filaments. In these systems, transitions in the network’s macroscopic properties (e.g. fluid-to gel-like (reviewed in [33]) depend on actin cross-linkers creating actin network connectivity so that myosin-II motor generated forces are transmitted some distance within the network beyond where force originates. Phase transitions in motor-driven transport have been observed in these network systems as well. For example, some studies have found that trajectories of myosin-II filaments transition from directed motion to becoming trapped in rings of actin filaments as the amount of cross-linking protein is increased [44, 45]. Other studies have found that with enhanced actin filament connectivity due to the addition of cross-linking proteins, directed myosin-II filament motion transitions to being stationary, as the motors undergo force-dependent stalling while engaged with separate actin filaments [46]. Although our liposome transport model similarly identifies actin filaments as vertices in a network, edges connecting these vertices are not physical cross-links as in the studies described above, but rather locations where a tug of war might occur as myoVa motors on the liposome surface engage separate actin filaments. Thus, in our model percolation does not underlie a phase transition in the mechanical properties of the actin network, but instead underlies the phase transition in the transport properties of the myoVa-liposome system. This phase transition involves both changes in motor trajectories, as they go from directed on subcritical networks to random walks on supercritical networks and, concurrently, a sudden increase in the number of stationary liposomes as tug of wars become much more probable. The existence of phase transitions in transport has implications in the regulation of intracellular transport, i.e., small changes in actin density or liposome diameter can have profound effects on motor-driven cargo transport.

### Actin network form and function

In cells, both spatial and temporal variations in the cytoskeletal actin density, may provide a mechanism to control the functional role that the actin network adopts (i.e., highway, sieve, or barrier; Fig. 8) through its impact on vesicular transport generated by the myoVa motors on the cargo surface. Therefore, the cell might exploit actin network transitions between subcritical and supercritical actin densities as a means of switching actin network functional roles thereby dictating cargo transport outcomes. For example, on sparse, subcritical actin networks, regardless of cargo size, the actin network would behave as a highway for directed and potentially targeted cargo delivery (Fig. 8). However, the percolation threshold for a given actin network does vary inversely with cargo radius, so that larger liposomes experience a transport phase transition at lower actin densities. Thus, a given actin network could act as a transport”sieve” (Fig. 8), whereby larger liposomes would get frequently caught up in tug of wars, while smaller liposomes would move relatively unencumbered to their targets along actin filaments (see SM for an example). Finally, at supercritical actin densities, the actin network may serve as a “barrier” (Fig. 8) as cargo and their myoVa motors transition predominantly to the tug of war state (Figs. 3, 7B). This condition has been described for insulin granules that are believed to exist in a readily releasable pool near the cell membrane within the dense cortical actin network [47, 48]. This barrier to granule delivery and docking to the cell membrane may be eliminated upon glucose stimulation by the regulated activity of actin severing (e.g. cofilin) and branch-forming (e.g. N-WASP/Arp2/3) proteins, creating a more favorable, subcritical density of actin filaments for myoVa granule delivery. This transition from myoVa being a tether to transporter further emphasizes the role that the actin filament network and it dynamic structure plays in cargo transport.

**Figure 8:**
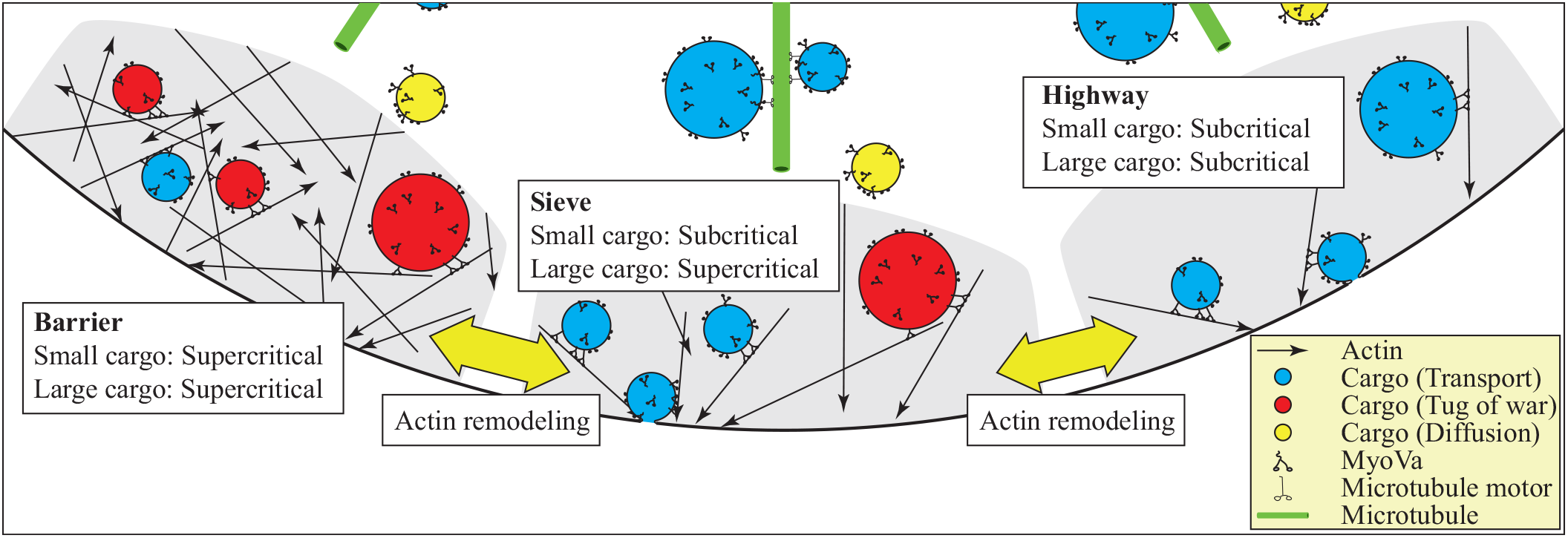
By varying actin density, cells can switch the functional role of the actin network to regulate intracellular transport.

In the future, building further complexity *in vitro* will require understanding of how cargo decorated with both actin-and microtubule-based motors share their transport duties in mixed cytoskeletal networks composed of both actin filaments and microtubules. Nevertheless, the modeling approach outlined here will allow these transport systems to be efficiently and accurately described over physiologically relevant spatial and temporal time scales.

## Materials and Methods

### Coarse-grained model

We developed a coarse-grained version of our experimentally-validated model for the transport of fluid, 350nm (radius *r*_*L*_ = 175nm) liposomes by a team of 10 myoVa motors through 3D actin networks [19, 20]. This coarse-grained model does not keep track of the myoVa motors on a liposome and their force and motion generation; instead at any given time, the liposome is assigned to one of three states: 1) transport, where one or motors are engaged with a single actin filament; 2) diffusion, where no motors are engaged with an actin filament, or; 3) tug of war, where some motors are engaged with one actin filament and other motors are engaged with a second actin filament. When in the transport state, the liposome moves along an actin filament towards the filament plus-end with a 36nm step 78% of the time, and with a short 31nm step 22% of the time. Due to the helical arrangement of binding sites on the actin filament with a repeat distance of 36nm, the liposome moves along an actin filament in a spiral path with an average period similar to experimental measurements [17,19]. When the liposome encounters a second actin filament within reach of the motors (*r*_*B*_ = 210nm from the liposome center, a parameter of the coarse-grained model determined from fitting simulations of our detailed model, and includes the liposome radius *r*_*L*_ = 175nm and the motors’ effective reach, see SM), the liposome may enter the tug of war state, where it becomes stationary. The tug of war is resolved when one of the motor teams generates a greater force allowing the liposome to enter the transport state on the actin filament associated with the winning team. A tug of war can also resolve by all motors detaching from actin and the liposome entering the diffusion state. In the diffusion state, the position of the liposome is determined at time intervals of Δ*t* = 0.5ms. The liposome takes a random step in (*x, y, z*) from a Gaussian distribution with standard deviation 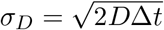 (for the radius *r*_*L*_ = 175nm liposomes, with diffusion constant *D* = 1.3*µ*m^2^*/*s, this is *σ*_*D*_ = 36nm). From the diffusion state, the liposome can enter the transport state when it collides with an actin filament. The small time step ensures that the simulation does not miss these collisions when they occur.

Transitions between states, and stepping are determined by rate constants that depend on the proximity of actin filaments to the liposome. Whereas the tug of war duration is dependent on the position and polarity of the two actin filaments with which the motors are engaged. These state transitions are determined with a Monte-Carlo method, using a modified version of the Gillespie algorithm (see Supplementary Material, SM, for details). The rate constants governing the state transitions, and the reach of the motors (*r*_*B*_), were determined by fitting the coarse grained model to thousands of simulations of our more detailed model [19, 20] (see SM for the fits). Since the Gillespie algorithm takes time steps of variable size and our simulation of diffusing liposomes takes constant time steps, the algorithm records liposome position and state at the constant time step and, when a state transition occurs between time steps, the new state is updated at the following time step (see SM for details).

### Liposome transport through a random actin network

We used this model to investigate how the density of actin filaments placed randomly in a network bounded by a 20*µ*m sphere affects 350nm (radius, *r*_*L*_ = 175nm) liposome transport. Each liposome starts at the network center bound to an actin filament. Therefore, one of the actin filaments in the network was constrained to pass through the center of the network sphere. For ease in modeling, the remainder of the actin filaments are placed randomly inside a 10 × 10 × 10*µ*m cube within the sphere. Their placement is accomplished by randomly selecting two points in the cube. An actin filament then connects those two points and extends in both directions to the network boundary. The actin filament polarity is assigned as the actin minus-end being nearer the first point and the plus-end nearer the second point.

For each actin network, we simulated the motion of 5,000 liposomes. Each simulated liposome trajectory concludes when either the liposome reaches the network boundary or 100 seconds elapse, whichever happens sooner. We performed these simulations on actin networks of 3, 6, 12, 25, 50, 100, 200, 400 and 800 filaments, covering roughly three orders of magnitude. For the networks with fewer filaments (3, 6, 12, 25, 50, 100), which typically gave more variable results, we examined *n* = 30 randomly generated networks. For the networks with more filaments (200, 400, 800), which gave more consistent results and were more computationally expensive, we examined fewer networks (*n* = 15, 8, and 5, respectively).

### Analysis of liposome trajectories

The output of each simulation was: 1) the 3D position and model state of each liposome at 0.0005s intervals over ≤ 100 seconds; 2) the position and polarity of each actin filament. The position and state matrices being at least 5,000 liposomes by 200,000 time points were too big to store and analyze easily. Therefore, after each simulation, matrices were compressed by only keeping every hundredth entry, equivalent to a time-step of 0.05s. In this way, we can still take a sufficiently small time step to ensure we don’t miss collisions between liposomes and actin, yet get simulated data with time steps similar to experimental measurements.

We measured four properties of each liposome trajectory: 1) average model state; 2) average speed; 3) mean squared displacement (MSD) scaling exponent for the entire trajectory, and; 4) standard deviation of the MSD scaling exponent calculated over 1s between *t* = 9 and *t* = 10s of the trajectory. These quantities were calculated as follows. The average model state for a given trajectory was determined as the proportion of simulation time that the liposome spent in each of the three states. For the 5,000 liposomes trajectories, the population average state is the average of these probabilities for the individual liposome trajectories. Average speed was the distance between the final and initial position of the liposome, divided by total simulation time. MSD scaling exponent was calculated by determining the mean squared displacement at a range of times, Δ*T*, from the time step 0.05s to half the total simulation time, and then calculating the best fit slope of MSD vs Δ*T* on a log-log scale. To calculate MSD scaling exponent for the entire trajectory, we performed the above calculation on the entire trajectory. For the standard deviation of the MSD scaling exponent, we performed the above calculation between *t* = 9 and *t* = 10s and obtained the standard deviation of this value for all 5,000 liposomes.

### Number of hit targets

As a measure of global transport within a given actin network and whether or not such transport could be considered “targeted”, we uniformly distributed 100 2*µ*m-diameter spherical targets on the network boundary sphere (see SM for details). A target was hit when any part of the liposome surface came into contact with a target. The number of targets hit provides a metric of the spread in the trajectories: if few targets are hit, then the trajectories are all approximately in the same direction; if many targets are hit, then the trajectories go in many different directions.

### Accessible actin

The number of accessible actin filaments in a given actin network is the set of potential filaments that can be reached by liposomes as they move from their starting point at the center of the network to the network boundary without detaching from actin (the algorithm, including validation, is given in the SM). This set determination encompasses the multitude of paths taken as a result of filament switching following a tug of war and that all paths must be in the direction of the actin plus-end. Therefore, not every actin filament will be in the set of accessible actin if access to an intersecting filament requires the liposome to travel towards the minus-end of a filament, which cannot occur.

### Simulations with larger liposomes

We performed a series of simulations as described above but with 5-fold larger, 1750nm diameter (*r*_*L*_ = 875nm), liposomes. The liposome diffusion constant was suitably adjusted (*D* = 0.26*µ*m^2^*/*s, giving a diffusion step size of *σ*_*D*_ = 16nm). Additionally, because the parameters of the coarse-grained model change with liposome size, we determined the rate constants governing the model state transitions and the reach of the motors (*r*_*B*_), by fitting the coarse grained model to thousands of simulations of our more detailed model [19, 20] in the same way as for the smaller liposomes (see SM for the fits).

For the simulations with larger liposomes, we constructed actin networks in the same way as for the smaller liposomes and also simulated the motion of 5,000 liposomes. However, we performed the simulations on actin networks of 3, 6, 12, 25, 50 and 100 filaments, covering roughly two orders of magnitude, since the simulations were generally more computationally expensive and to ensure that the pore sizes of the networks were larger than the liposome diameter (see SM). For the networks with fewer filaments (3, 6, 12), which typically gave more variable results, we examined *n* = 30 randomly generated networks. For the networks with more filaments (25, 50, 100), which gave more consistent results and were more computationally expensive, we examined fewer networks (*n* = 20, 10, and 5, respectively).

### Simulations with initially diffusing liposomes

We ran simulations with both small (350nm, *r*_*L*_ = 175nm) and large (1750nm, *r*_*L*_ = 875nm) liposomes initially detached from actin and therefore in the diffusion state. Simulations were performed identically as for the case where liposomes start attached to an actin filament, except that we ensured that no actin filament passed through the liposome’s initial position. Actin networks were also constructed identically, except we did not constrain one actin filament to pass through the center of the network.

## Acknowledgments

The authors are grateful to Brandon Bensel for helpful discussion and feedback on an early version of the manuscript. This work was supported by grants from the National Institutes of Health R01GM094229 and R35GM141743 (to D.M.W.).

## Notes

### Competing Interest Statement

The authors have declared no competing interest.

