## Supplementary Material for "Modeling myosin Va liposome transport through actin filament networks reveals a percolation threshold that modulates transport properties"

S. Walcott and D. M. Warshaw

#### 1 The coarse-grained model

To construct a simplified model of molecular motors transporting cargo through an actin network, we must capture the essential features of this process. As a team of myosin motors transports their cargo through an actin network, they alternate between interacting with a single actin filament and, potentially, interacting with a second actin filament[1]. At these actin "intersections" a tug of war between motor teams can arise, and the cargo can 1) be transported on the new actin filament; 2) be transported on the original actin filament; or 3) detach from actin. The relative position of the two actin filaments affects both the duration of the tug of war, and the probability of each of the three outcomes. Our model [1,2], henceforth called the Detailed Model, successfully predicts these quantities for the transport of 350nm diameter fluid liposomes transported by $\sim 10$ myosin Va molecular motors. To develop a simplified and more computationally efficient model, we performed a systematic analysis of the behavior of myosin-mediated transport through actin intersections. We thereby generated a simulated data set on which we trained (i.e. tested the assumptions and tuned the parameters of) our simpler model.

##### 1.1 Systematic analysis of pairwise intersections

The relative position of any two actin filaments can be defined by two variables, their minimum separation ($z$) and an angle ($\theta$).[2] Thus, as a first step toward developing more efficient simulation methods, we performed a systematic analysis of intersections, performing 50 simulations of the detailed model at $z = 0$, 100, 200, 300, and 400nm, at $\theta = 2\pi/16, 3\pi/16, \ldots, 14\pi/16$. A total of 3,250 individual simulations, under 65 different conditions.

For a given pair of filaments, there is an interaction zone, which describes the set of points where the liposome is positioned such that motors can simultaneously interact with both actin filaments (Fig. S1A). For each of the 65 different simulation conditions, we determined the number of simulations where the liposome 1) exited the interaction zone along the original filament (straight outcome); 2) exited the interaction zone along the other filament (turn outcome); or 3) detached within the interaction zone (terminate outcome). The turn and terminate outcomes become more likely as the filaments become more parallel (i.e. with plus ends aligned) and antiparallel (i.e. with plus ends pointing in opposite directions), because the size of the interaction zone increases under these conditions, allowing multiple tug of wars. Similarly, as filament separation increases, so too does the likelihood of the straight outcome (Fig. S1B).

[1] according to our model, concurrent interactions with three filaments are rare, [2]

[2] Suppose that $\boldsymbol{x}_i^s$ and $\boldsymbol{x}_i^e$ are vectors to the start (minus end) and end (plus end), respectively, of actin filaments $i = 1$ and 2. We can then find unit vectors pointing along each filament, in the direction of their polarity (i.e., toward their plus ends) $\hat{\boldsymbol{e}}_i^r = (\boldsymbol{x}_i^e - \boldsymbol{x}_i^s)/||\boldsymbol{x}_i^e - \boldsymbol{x}_i^s||$. Then, we can find a unit vector perpendicular to both filaments by $\hat{\boldsymbol{e}}^p = (\hat{\boldsymbol{e}}_1^p \times \hat{\boldsymbol{e}}_2^p)/||\hat{\boldsymbol{e}}_1^p \times \hat{\boldsymbol{e}}_2^p||$. Then, $z = |(\boldsymbol{x}_1^s - \boldsymbol{x}_2^s) \cdot \hat{\boldsymbol{e}}^p|$, and $\theta = \cos^{-1}(\hat{\boldsymbol{e}}_1^r \cdot \hat{\boldsymbol{e}}_2^r)$.

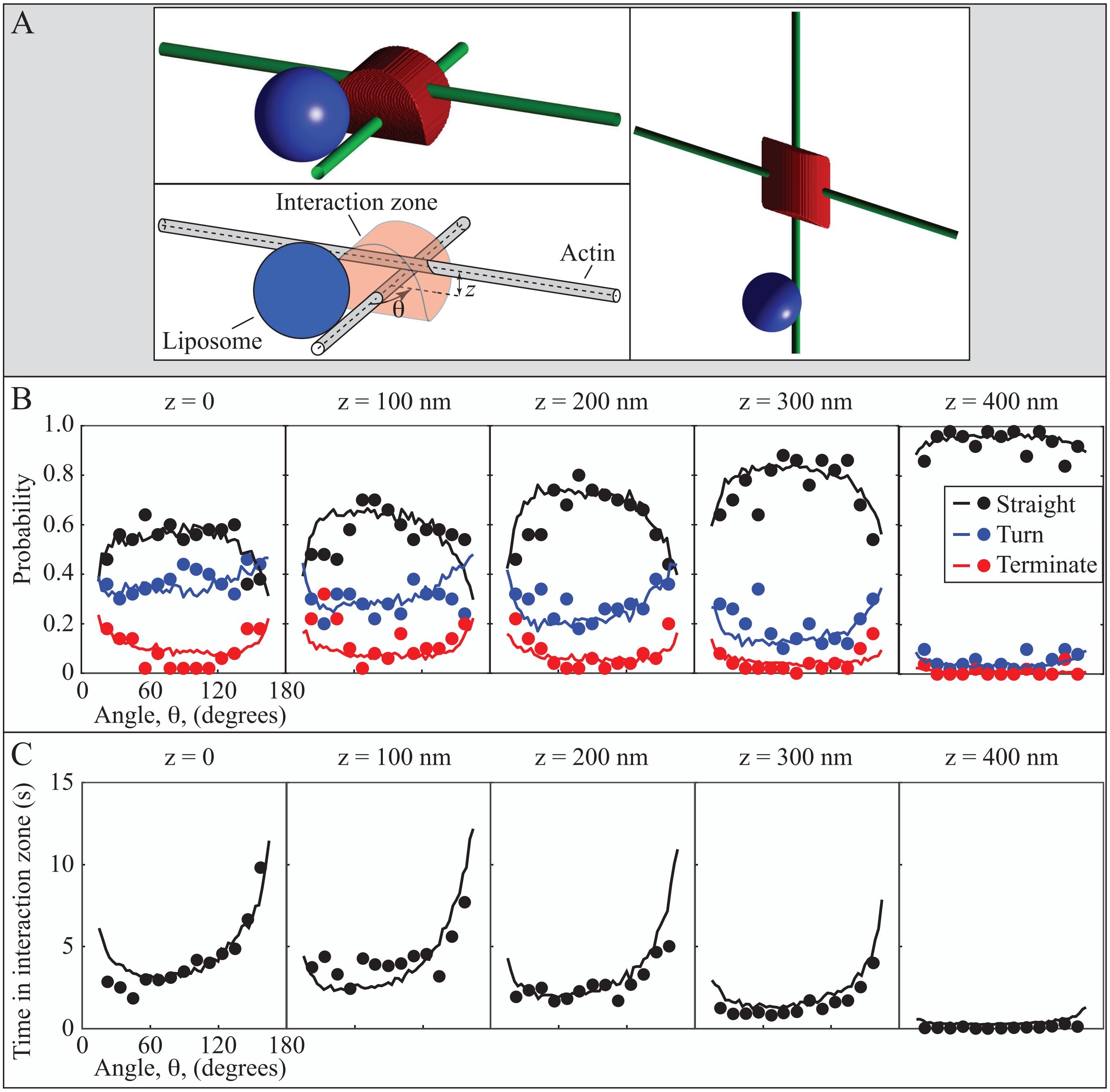


Figure S1: A coarse-grained model captures the behavior of 350nm diameter fluid liposomes transported by 10 myosin Va motors through actin intersections. A. Schematic of a liposome approaching an actin intersection with separation $z$ and angle $\theta$. The interaction zone, where myosin motors on the liposome can interact with both actin filaments, is shown in red. B. Outcomes (the liposome passing straight through, turning at, or terminating at the actin intersection) for intersections of different angle ($\theta$) and separation ($z$). See Movie S15 for one example simulation. C. The time the liposome spends in the interaction zone for intersections of different angle ($\theta$) and separation ($z$). In B and C, dots are simulations of full model, $N = 50$, and lines are simulations of the coarse-grained model, $N = 1{,}000$. The scaling on each subplot is identical, but the horizontal axis is only labeled for the $z = 0$nm plot (left).

Besides the outcomes of each simulated liposome trajectory, we also determined the time that each trajectory spent in the interaction zone. As the filaments become more parallel, the size of the interaction zone increases, and therefore the time spent in it naturally increases. Additionally, there are more opportunities for tug of wars, which would also contribute to increasing the time in the interaction zone. However, there is a strong asymmetry with angle – that is, liposomes move relatively efficiently through the interaction zone of more-or-less parallel filaments, and slowly through the interaction zone of more-or-less antiparallel filaments, even though they are the same size (Fig. S1C).

#### 1.2 The reduced model

The goal of our reduced model is to capture the behavior of the detailed model, while being both simpler and more computationally efficient. At the same time, we want to provide a means to connect our reduced model to the detailed model, so that we minimize the loss of information with the model reduction. To achieve these tasks, we develop a mechanistic model containing a few parameters representing course-grained effects at the molecular level. We then determine these parameters by fitting our simulation results. In this way, should parameters of the detailed model change, we can re-run these simulations and determine the corresponding changes to the parameters of the reduced model (see Section 1.3). We now describe the reduced model, and show the results of our fits to the simulations.

##### 1.2.1 Model formulation

In the reduced model, we do not explicitly track the motors on the liposome. The liposome's position on an actin filament is defined by the distance from the minus end of that filament, $s$, and an azimuthal angle $\phi$. Then, the liposome steps forward at a rate $k_{step} = 10\text{s}^{-1}$ and, with probability $p_s = 0.22$ takes a short step (31nm, with a change in $\phi = \Delta\phi = \pi/7$ – given that the motor makes a full rotation for every 14 short steps) and with probability $1 - p_s$ takes a step (36nm, with no change in $\phi$) along actin's repeat distance. When a second actin filament comes within the reach of the liposome, $r_B$, a tug-of-war may be initiated, and occurs at rate $k_{tw}$. Each tug of war, once started, results in a termination with probability $P_{term}$, a turn with probability $P_{turn}$ and a straight outcome with probability $P_{straight} = 1 - P_{term} - P_{turn}$ (Fig. S2). This tug of war is resolved with a rate constant, $k_R$, which depends on the relative angle between the two filaments ($\theta$):

$$\frac{1}{k_R} = t_R = t_0 \exp\left(c \cdot \cos\left(\frac{\pi - \theta}{2}\right)\right)$$

This model then has the following unknown parameters, $t_0$, $c$, $P_{turn}$, $P_{straight}$, $k_{tw}$. In addition, the parameter $r_B$, the reach of the motors, must be between the radius of the liposome $r_L = 175$nm and that radius plus the length of the motor $r_L + \ell = 175 + 50 = 225$nm. We can adjust these parameters to fit the simulations of the full model. The values $P_{turn} = 0.4$, $P_{term} = 0.066$, $k_{tw} = 2\text{s}^{-1}$, $t_0 = 1$s, $r_B = 210$nm and $c = 1.1$ give reasonably good agreement with the simulations of the detailed model (Fig. S1B,C). This reduced model is roughly 10,000 times more efficient than the detailed model – e.g., the 3,250 simulations that take 1-2 weeks of computer time with the full model can be accomplished in minutes. The computational efficiency of this model allows us to perform the millions of simulations described in the main text.

#### 1.3 Larger liposomes

Besides simulating the motion of liposomes of radius $r_L = 175$nm, we also simulated the motion of liposomes of radius $r_L = 875$nm. Since the larger liposome has five times the radius of the smaller, it has 25 times the surface area. Thus, to keep the same motor density, we must increase the number of myoVa motors from 10 to 250 in the detailed model. As with the smaller liposomes, we performed a systematic analysis of intersections, performing 50 simulations of the detailed model at $z$ = 0, 100, 200, 300, and 400nm, at $\theta = 2\pi/16, 4\pi/16, \ldots, 14\pi/16$. A total of 1,750 individual simulations, under 35 different conditions. Adjusting the parameters of the reduced model to fit the simulations of the full model, we get reasonably

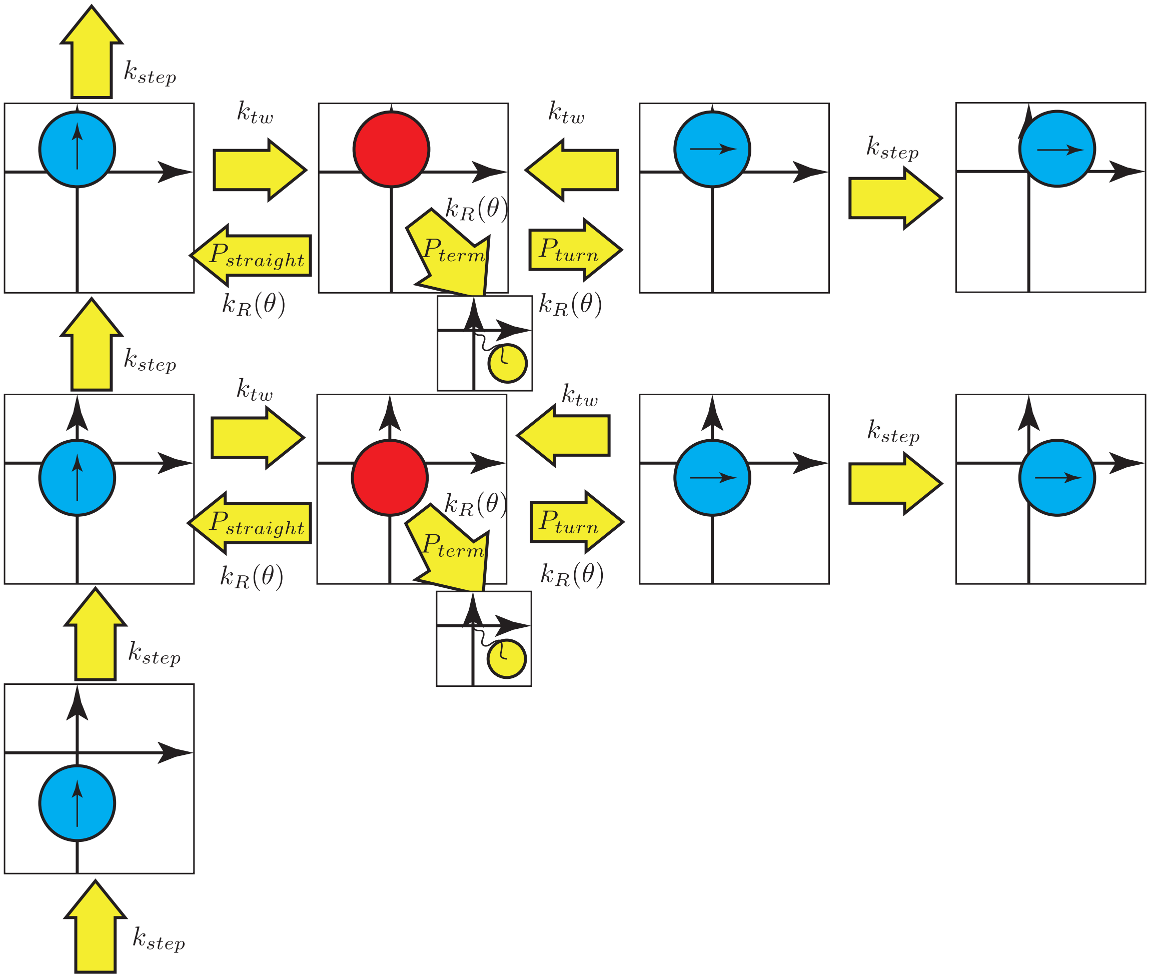


Figure S2: Schematic of the kinetics of the reduced model. Starting in the lower left corner, the liposome in the transport state (blue) steps forward along its associated actin filament at a rate $k_{step}$. When within reach of a second actin filament, the liposome may enter the tug of war state (red) with rate $k_{tw}$. The tug of war is resolved at rate $k_R(\theta)$ that depends on the relative angle of the two actin filaments, $\theta$. There are three possible resolutions to the tug of war: the liposome can continue along the original actin filament (with probability $P_{straight}$), the liposome can switch to the second actin filament (with probability $P_{turn}$), or the liposome can detach from both actin filaments and enter the diffusion state (yellow, with probability $P_{term}$).

good agreement (Fig. S3A,B). Model parameters are as follows: $P_{turn} = 0.4$, $P_{term} = 0.03$, $k_{tw} = 1\text{s}^{-1}$, $t_0 = 1\text{s}$, $r_B = 910\text{nm}$ and $c = 1.1$.

#### 2 Simulation details

As discussed above (Section 1.2.1), in the reduced model, we keep track of the state of a liposome. That is, a liposome can either be in 1) a transport state, 2) a tug of war state, or 3) a diffusive state. Transitions between these states are given by the rate constants described in that section. To implement this, we use a modified version of the Gillespie algorithm, as in Lombardo et al. 2017, 2019. However, since liposomes may diffuse in these simulations, we must modify the simulation procedure to account for diffusive motion.

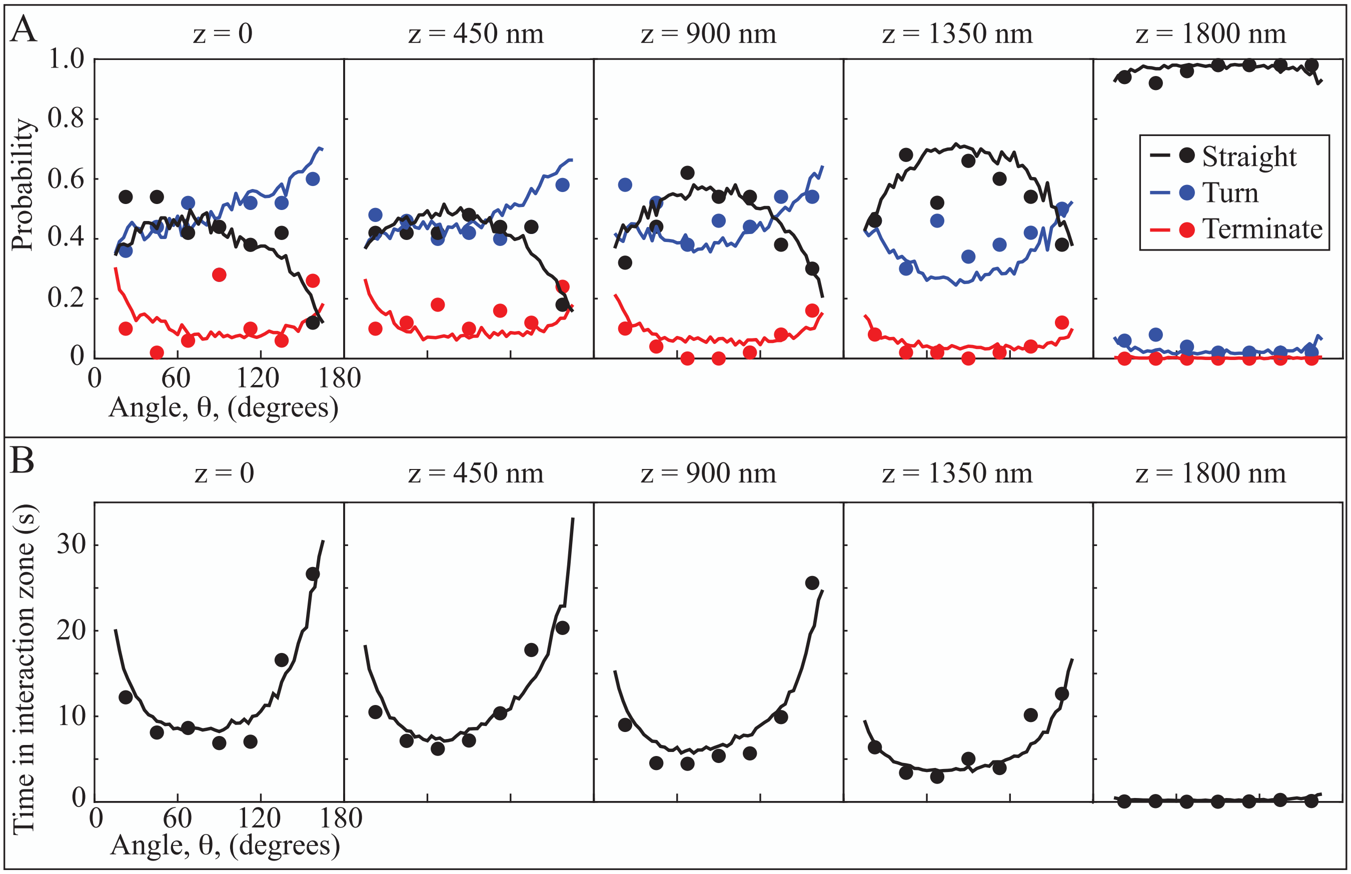


Figure S3: A coarse-grained model captures the behavior of 1,750nm diameter fluid liposomes transported by 250 myosin Va motors through actin intersections. A. Outcomes (the liposome passing straight through, turning at, or terminating at the actin intersection) for intersections of different angle ($\theta$) and separation ($z$). B. The time the liposome spends in the interaction zone for intersections of different angle ($\theta$) and separation ($z$). In A and B, dots are simulations of full model, $N = 50$, and lines are simulations of the coarse-grained model, $N = 1{,}000$. The scaling on each subplot is identical, but the horizontal axis is only labeled for the $z = 0$nm plot (left).

#### 2.1 Diffusion

To simulate diffusion, we must first calculate the diffusion constant for the liposome. We assume Stokes' drag on a sphere and calculate diffusion constant from Einstein's relation, $D = \frac{k_B T}{6\pi\eta r_L}$, where $k_B$ is Boltzmann's constant, $T$ is the absolute temperature, $\eta$ is the dynamic viscosity of water at the appropriate temperature, and $r_L$ is the radius of the liposome. We assume that the system is immersed in water at 25°C. Note that the viscosity in cells larger than this, and temperature is also higher. For liposomes with $r_L = 175$nm, we calculate $D = 1.3 \cdot 10^6 \text{nm}^2/\text{s}$, and with $r_L = 875$nm, we calculate $D = 0.26 \cdot 10^6 \text{nm}^2/\text{s}$.

With the diffusion constant, we can update the position of a diffusing liposome in the following way. If the position of the liposome at time $t$ is $\boldsymbol{x}(t)$, its position a short time $\Delta t$ later is

$$\boldsymbol{x}(t + \Delta t) = \boldsymbol{x}(t) + \boldsymbol{X}(0, \sqrt{2D\Delta t})$$

where $\boldsymbol{X}$ is a vector of three Gaussian distributed random variables with mean 0 and standard deviation $\sqrt{2D\Delta t}$. At each time step, we check whether the liposome collides with an actin filament. If so, we assume that one of the motors engages with that filament. If not, the liposome continues diffusing.

Simulating diffusion requires a choice of time step $\Delta t$. It is important to pick a sufficiently small $\Delta t$ so that we can detect each time that the liposome collides with an actin filament. This requires that the expected step size, $\sqrt{2D\Delta t}$, be much less than the radius of the liposome. We picked $\Delta t = 5 \cdot 10^{-4}$s, which gives an expected step size of 36nm for the smaller liposomes ($r_L = 175$nm), and 16nm for the larger liposomes ($r_L = 875$nm). With this time step, we do likely miss some collisions where the liposome just grazes an actin filament, but we expect that we capture the vast majority of collisions.

##### 2.2 Simulating chemical reactions and diffusion

The Gillespie algorithm takes time steps dictated by rate constants and random numbers drawn from the appropriate distribution. The diffusion simulation takes a constant time step. Thus, to implement the simulations, we separately keep track of all liposomes that are diffusing and all liposomes that are undergoing chemical reactions (i.e., that are in the transport or tug of war state). For those undergoing chemical reactions, we start by calculating a vector of reaction times $\boldsymbol{t}_c$, using the Gillespie algorithm. We then define a vector of times from 0 to 100s, spaced by $\Delta t = 5 \cdot 10^{-4}$s.

At time $t = 0$, the state of each liposome is known. In particular, we performed two different simulations, one where all liposomes start detached from the actin network and one where all liposomes start attached to the actin in the transport state. To calculate the state and position of all liposomes at time $\Delta t$, we find all of the entries of $\boldsymbol{t}_c$ that are less than $\Delta t$. These are the chemical reactions that occur within that time step. These chemical reactions include stepping, tug of war initiation, tug of war termination and detachment. These reactions are updated and the position of these liposomes updated, until all of the entries of $\mathbf{t}_c$ are greater than $\Delta t$. We then update the position of all diffusing liposomes, and record the state and position of all liposomes at time $\Delta t$. This process is iterated until we reach the end of the simulation, $t = 100$s.

One problem is that liposomes collide with actin filaments and then switch from the diffusion to the transport state. Similarly, liposomes in tug of wars may detach from actin and enter the diffusion state. These, in general occur at times in between the entries of our vector of times at which we record the state of the liposomes. For simplicity, we assume that these occur at the time step previous to the detachment/attachment event. This introduces an error on the order of $\Delta t < 5 \cdot 10^{-4}$s which, given that reactions occur on time scales of $\sim 1$s, is small.

##### 2.3 Distributing targets

To assess how targeted transport is on a given actin network, we distributed 100, 1$\mu$m radius targets evenly over the surface of the sphere that represents a transport distance of 10$\mu$m. To do so, we wrote a simulation where the targets, which are constrained to move on the surface of the sphere, repel each other. We then ran this simulation until it approached steady state.

In the simulation, each target experiences a force from all the other targets. The force on the $i^{th}$ target, $\boldsymbol{F}_i^T$ is

$$\boldsymbol{F}_i^T = \sum_{j \neq i} \frac{1}{(\boldsymbol{x}_i - \boldsymbol{x}_j)^2} \hat{\boldsymbol{e}}$$

where $\boldsymbol{x}_i$ (or $\boldsymbol{x}_j$) is the position of the $i^{th}$ (or $j^{th}$) target, and $\hat{\boldsymbol{e}}$ is a unit vector that points from $\boldsymbol{x}_j$ to $\boldsymbol{x}_i$.

In addition to this force, each target experiences a viscous drag proportional to its velocity, $\boldsymbol{F}_i^D = \gamma \frac{d\boldsymbol{x}_i}{dt}$. Then, assuming the targets move in the inertialess regime, the equation of motion are

$$\boldsymbol{F}_i^T + \boldsymbol{F}_i^D = \mathbf{0}$$

which, with the definition of $\boldsymbol{F}_i^D$, we can rearrange to

$$\frac{d\boldsymbol{x}_i}{dt} = -\frac{1}{\gamma} \boldsymbol{F}_i^T$$

We then numerically solved this equation with a forward Euler scheme. At each time step, the target moves a little bit off the sphere, so we projected its position back onto the sphere. We chose $\gamma = 1$, a time step of $\Delta t = 1 \cdot 10^{-3}$, and ran the simulation for 5,000 time steps.

### 3 Analysis of actin networks

When liposomes are transported by myoVa motors along an actin filament, they may or may not encounter another actin filament with which the myoVa motors may interact. If they do, there are then two potential paths available if the motors do not detach: the motors might continue along the original actin filament, or they might switch to a new filament. As the liposome continues past this first intersection, on whichever path, it may encounter another actin filament and give rise to other potential paths. We performed a series of simulations to demonstrate that there is a critical number of actin filaments, $N_c$, above which there is an abrupt increase in the scaling of average number of paths with number of actin filaments. Here, we describe the simulations that allow us to determine $N_c$.

The model neglects steric interactions between the liposomes and the actin network. This is a reasonable assumption, provided that the diameter of the liposome is less than the pore size of the actin network. We therefore estimated the pore size of our actin networks, and show that it is much greater than the liposome diameter at $N_c$. Here, we present this analysis, and also a more precise definition of pore size.

#### 3.1 Transport path algorithm to determine the number of accessible actin

Given a network of actin filaments, it is not so straightforward to find all of the actin filaments that may be reached by a liposome being transported by myoVa. Complexity arises because one must account for the fact that transport occurs toward the plus end of each actin filament; therefore, even though two actin filaments may be close enough for a liposome to simultaneously interact with both, the liposome may never reach this intersection because it occurs toward the minus end of the actin filament on which the motors transport the liposome.

Here is a description of the algorithm we use to identify all actin filaments along which transport may occur, which we term the number of accessible actin filaments:

1. We first calculate a matrix of the minimal distance between each pair of filaments, $\mathbf{D}$.

2. We identify all potential intersections, where $D_{ij} \leq 2r_B$, where $r_B$ is the radius of the liposome plus the effective reach of the motors (a fitting parameter of the course-grained model).

3. We calculate where this intersection occurs on the actin filament, relative to where the minus end of the actin filament intersects the edge of the spherical domain.

4. We discard all potential intersections that occur outside the spherical domain. This introduces a small error, since this minimal distance may occur outside of the domain, yet the filaments may still approach within $2r_B$.

5. Starting from the initial position of the liposomes (halfway along actin filament 1), we find all intersections between that point and the end of filament 1. For each of these intersection, we record the number of the actin filament and the relative position at which the intersection occurs.

6. We repeat this process for each of the new filaments, finding all filaments that intersect it between the intersection point and its end.

7. The process finishes when we no longer identify new actin filaments.

8. We then remove redundancies, to find all unique actin filaments that can be reached by a transported liposome, the accessible actin filaments.

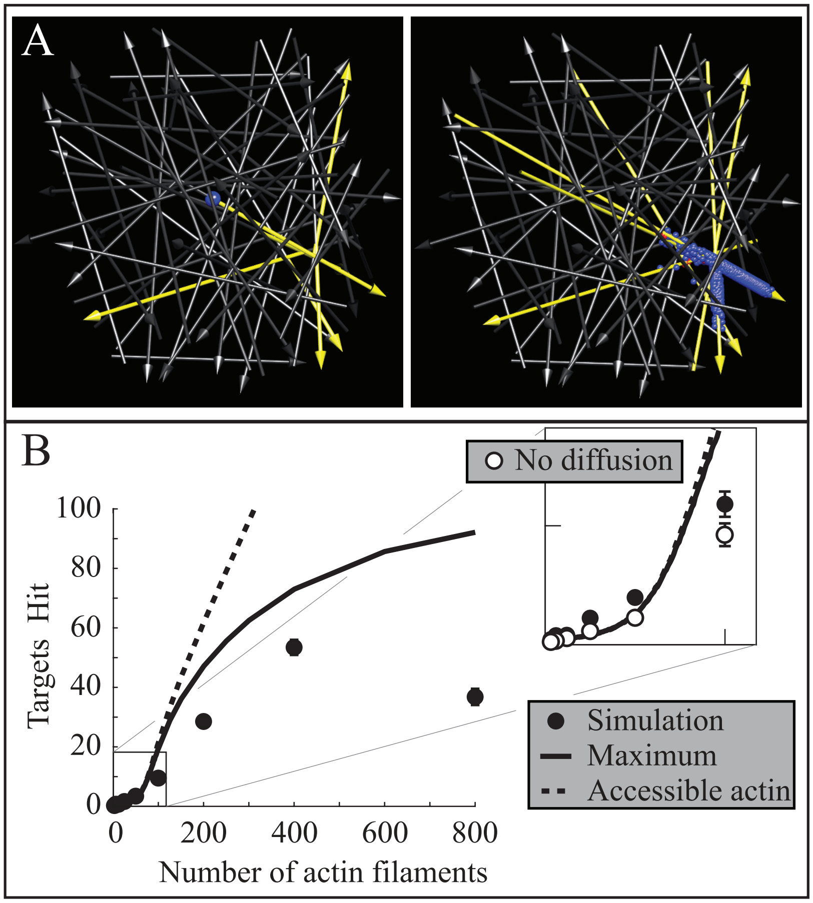


Figure S4: A. The path following algorithm identifies six accessible actin filaments on an actin network of 50 filaments (left, from Fig. 3 of main text, yellow shows possible paths along the accessible filaments). Liposomes that do not detach from the actin network interact with the same six filaments (see supplementary movie S8 for the full simulation). B. The measured number of hit targets is comparable to the maximum number of hit targets and the accessible actin multiplied by 0.35, the probability of an actin filament terminating at a target. Error bars (mostly obscured by symbols) show SEM.

##### 3.2 Validation of the algorithm

To ensure that the algorithm successfully identifies accessible actin filaments, we performed two different simulations. In the first, we calculated the accessible actin on two different networks, one with 50 actin filaments (a subcritical network, see Section 2.5 of the main text for a definition of "subcritical") and one with 100 actin filaments (a supercritical network, see Section 2.5 of the main text for a definition of "supercritical"). The two networks and the accessible actin are shown in Fig. 4 of the main text. We then ran simulations of 2000 liposomes being transported through those networks. Once complete, we visualized the motion of only those liposomes that remained bound to actin for the entirety of the simulation, and did not enter the diffusive state (1972 liposomes on the 50 filament network; 1934 liposomes on the 100 filament network). If our algorithm worked correctly, then the visualized liposomes should only interact with accessible actin filaments. This is indeed what we observe. For example, in the network with 50 filaments, there are six accessible actin filaments and, at one point in the movie, liposomes interact with all six (Fig. S4A) and at no point in the movie do they interact with any others (see movie S8 for the full simulation). For the network with 100 filaments, much more of the network is accessible (78% vs. 12%), and the liposomes do not interact with all accessible actin filaments (discussed in more detail below). Nevertheless, they exclusively interact with filaments that are identified as accessible (see movie S9 for the simulation, cf. Fig. 4 of the main text).

In the second set of simulations we performed, we calculated the number of accessible actin filaments for 2,000 different actin networks of sizes 1-125, 150, 200, 300, 400, 600 and 800 filaments. We then calculated the "maximum" number of targets that could be hit. To do so, we determined the number of targets that are within $r_B$ of the ends of these filaments. This identifies the maximum number of targets that that could be hit by liposomes that do not detach from the actin network. Since liposomes do detach from the actin network, this is not a true maximum, but is a good approximation because these detachment events are rare. Thus, if the calculation is correct, then we expect that on subcritical networks, this "maximum" is a slight underestimate. The situation changes on supercritical networks, where there is a sudden increase in the number of potential travel paths. Even with our simulations including a large number (5,000) of liposomes, not all of these potential paths are traveled by a liposome (this can be observed in movie S9). As a result, we expect the "maximum" number of targets we calculated to be an overestimate. Indeed, we observe these expected results (Fig. S4B). Further, on subcritical networks when we only consider liposomes that never enter the diffusion state (liposomes that diffuse make up $< 2\%$ of the total on these subcritical networks), the agreement between the calculated and observed number of hit targets is excellent (Fig. S4B, inset), giving further credence to our algorithm.

##### 3.3 Connection between accessible actin and hit targets

In Fig. 4A-E of the main text, we show that a variety of properties of liposome transport scale differently on subcritical and supercritical actin networks. This phase transition arises because, as discussed in the main text, the relatively sudden increase in transport paths that occurs on the switch from subcritical to supercritical networks results in an increase in tug of wars that hold the liposome in place, and a loss of directionality that occurs when liposomes switch filaments at the resolution of these tug of wars. Though this provides a qualitative understanding of the phase transition, it is challenging to provide a quantitative description for most of these properties. However, for the number of hit targets, we can provide a quantitative description.

On the subcritical actin networks where relatively few transport paths occur, the plus ends of the accessible actin filaments are relatively far apart. Therefore, since the probability of a given actin filament contacting a target is 0.35, the number of hit targets should be well-approximated by 0.35 times the amount of accessible actin. This argument assumes that each actin filament is independent of the others, an assumption that breaks down on dense actin networks with many transport paths. On such networks, the probability that two (or more) actin filaments terminate at the same target is not negligible, so the approximation greatly overestimates the number of hit targets. Nevertheless, for subcritical actin networks and through the phase transition, we expect that this approximation will be good. Indeed, we observe excellent agreement up to networks of around 70 actin filaments (dashed line, Fig. S4B), providing a quantitative description of the change in scaling in the number of hit targets.

##### 3.4 Estimating $N_c$ from mean accessible actin

When accessible actin undergoes a phase transition, we see a power law scaling of the form $\alpha N^{\beta}$ on subcritical networks (below a critical number of actin filaments, $N_c$), and a different scaling $a(N-N_c)^b$ on supercritical networks (above $N_c$, Fig. 4E fo the main text). The former fit should apply when $N$ is small, and become poor near $N_c$. Therefore, to we fit the number of accessible actin filaments on networks of 2-10 filaments, minimizing the $\chi^2$ error. The best-fit parameters were $\alpha = 0.908$ and $\beta = 0.142$.

The scaling $a(N - N_c)^b$, should apply above $N_c$; however, it should not apply too close to $N_c$, and it might not apply far from $N_c$. We therefore fit the number of accessible actin filaments on networks of 70-150 filaments, minimizing the $\chi^2$ error. Since we did not measure the number of accessible actin filaments on actin networks with 126-149 filaments, we interpolated between our measurements and their standard deviation to ensure an equal weighting of the measurements with network size. The best-fit parameters were $a = 2.61$ and $b = 0.862$ and $N_c = 59.7$. These best fit values have a modest dependence on the network sizes used for the fits, with $a$ ranging between 1 and 4, $b$ between 0.75 and 1, and $N_c$ between 55 and 65. We selected the range that gave, in our judgment, the most plausible fit.

##### 3.5 Estimating $N_c$ from distributions of accessible actin

In Fig. 6 of the main text, we show the results of another set of simulations in which we calculated the number of accessible actin filaments for 10,000 different actin networks of sizes 1-100, 125, 150, 200, 250, 300 and 400 filaments and then determined the proportion of the total actin that were accessible. To identify $N_c$, we calculated when the probability distribution of this proportion of actin switches from being uni- to bimodal. To do so, we had to accurately identify the mode of the distribution, which is straightforward everywhere except near $N_c$, where the distribution is transitioning from uni- to bimodal. There, since the distribution is relatively flat, the mode is particularly sensitive to noise. We employed two methods to minimize the influence of this noise.

In the first method to calculate the mode(s) of the distribution, we take a weighted average of the numbers of accessible actin that are within 90% of the maximum. For networks of $\sim 45$ filaments or less, this identifies the maximum at a single accessible actin filament. To find the second mode, when present, we discard all accessible actin filaments with fewer than 17 filaments, and then re-take a weighted average of the numbers of accessible actin that are within 90% of the maximum. When the distribution is unimodal, this identifies a mode near 18 filaments; when the distribution is bimodal, this identifies a mode that is larger than that. The switch from unimodal to bimodal occurs at $N_c = 58$ filaments. Different choices of the various cutoffs has a modest effect, but $N_c$ remains in the range 55-60. The cutoffs that we chose gave, in our judgment, the most accurate location of the mode(s).

In the second method to calculate the mode(s) of the distribution of accessible actin, for each actin network size, we fit the histograms of the proportion of accessible actin in the total network with a skew-normal and an exponential distribution. We then calculated the mode of the best fit skew-normal distributions (the exponential distribution has a mode of 1 accessible filament). This mode rapidly moves away from one accessible actin filament at $N_c = 58$ filaments, and agrees well with the mode calculated with the first method.

##### 3.6 Pore size

For a given 3D network of filaments in water, sufficiently small spherical particles can diffuse relatively unencumbered, while sufficiently large particles are trapped. The diameter of the largest particle that can pass through the network is the pore size of the network. The pore size depends on the density of the 3D network, details of how the network is arranged (whether the filaments are, e.g., randomly aligned or parallel), and the distance the particle is desired to travel.

We estimated the pore size of the actin networks used in our simulations, using the following algorithm:

1. A point is randomly selected within 10nm of the center of the 3D filament network

2. A sphere is grown from that point by a) drawing a sphere whose center is that point, and which is tangent to the nearest actin filament; b) increasing the diameter of the sphere until the sphere is tangent to a second actin filament, moving the sphere's center along a vector starting where the sphere touches the actin filament and ending at the center of the sphere; c) increasing the diameter

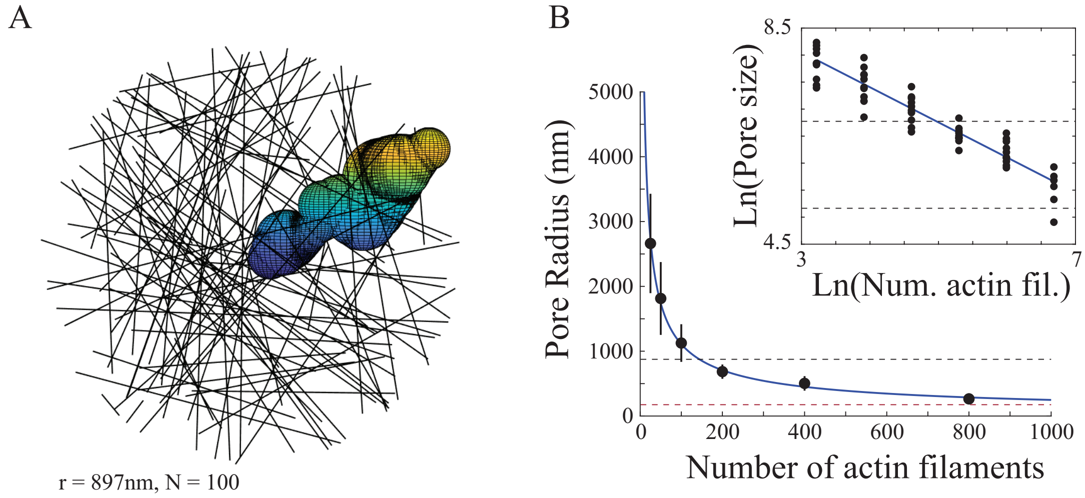


Figure S5: Pore size in random 3D actin meshes. A. An example calculation of a pore in a mesh of 100 actin filaments. The image shows a series of overlapping spheres, each of which is just tangent to three actin filaments. The pore radius, 897nm in this simulation, is the radius of the smallest sphere, ensuring that any smaller sphere could move from the center of the mesh to the boundary without touching an actin filament. See Movie S16 for a more detailed look at the pore calculation. B. Pore radius as a function of the number of actin filaments obeys a power law scaling. Simulation results (error bars show SD) are well-fit by a power law, and the simulation results are approximately linear when plotted on log-log axes (inset). Dashed lines show the radius of the smaller ($r_L = 175$nm) and larger ($r_L = 875$nm) liposomes.

of the sphere, moving its center along a vector orthogonal to both actin filaments it touches, until it is tangent to a third actin filament.

3. A point is selected in that sphere, toward the exterior of the network. Randomness is introduced in the size of this step (a uniformly distributed random variable, $w$, between 0 and 1 times 0.3 of the radius of the sphere) and also in the direction ($0.03(1 - w)$ times a random unit vector).

4. A sphere is grown at this new point, and the process repeats until the center of the sphere is outside 10$\mu$m from the center of the network, which is the travel distance used in our simulations.

5. It is determined whether all of the spheres overlap (spheres overlap if the center of one is located within one radius' distance of the other).

6. The smallest spheres are successively removed until removal causes the remaining spheres to no longer overlap.

7. A minimal set of overlapping spheres is found by removing spheres that are not necessary for the remaining spheres to overlap.

8. The radius of the smallest sphere in this minimal set defines the radius of a pore from the center of the mesh to a distance 10$\mu$m away.

For steps 5-8, we calculate the adjacency matrix for the spheres, a symmetric matrix $\boldsymbol{M}$ whose entry $m_{ij} = 1$ if spheres $i$ and $j$ overlap, and $m_{ij} = 0$ if not. Adjacency matrices arise in graph theory, the study of networks of vertices and edges. For an undirected graph, if the $i^{\text{th}}$ and $j^{\text{th}}$ vertices are connected with an edge, then $m_{ij} = 1$ and $m_{ij} = 0$ if not. The $n^{\text{th}}$ power of the adjacency matrix gives the number of $n$-step paths between the vertices. That is, if the $i, j$ entry of $\boldsymbol{M}^4$ is $m_{ij}^4 = 2$, then there are 2 distinct ways that one could travel from the $i^{\text{th}}$ to the $j^{\text{th}}$ vertex in 4 steps, where one moves from one vertex along an edge to another vertex each step (one can also remain at the same vertex in a step, since vertices are connected to themselves). So, given $n$ spheres from steps 1-4 of our algorithm, to determine whether a path exists from the sphere located at the center of the actin network (sphere 1) and the sphere whose center is outside of the network (sphere $n$), we look at the $1, n$ entry of $\boldsymbol{M}^{n-1}$. If $m_{1n}^{n-1} = 0$, then there is no path from the first to the $n^{\text{th}}$ sphere and so the spheres do not all overlap; if $m_{1n}^{n-1} \neq 0$, then the spheres do overlap.

This algorithm finds the radius of a pore through the mesh (Fig. S5A and Movie S16 shows an example). To estimate the size of the maximal pore, we ran this algorithm ten times on each mesh, and record the largest pore radius. Note that this is necessarily an underestimate, since our algorithm might not have found the largest pore allowed by a given network. For actin meshes of 25, 50, 100, 200, and 400 filaments, we ran this algorithm on ten different actin meshes; for actin meshes of 800 filaments, we ran the algorithm on six different actin meshes. The pore radius shows a power law scaling, that goes as the number of actin filaments to the $-2/3$ power (Fig. S5B).

From these simulations, we find that pore radius is larger than the radius of the liposome for all of our simulations of myoVa-mediated liposome transport. Importantly, at $N_c$ the pore radius is $\sim 1,750$nm for the 175nm radius liposomes, and $\sim 5,000$nm for the 875nm radius liposomes. Therefore, we conclude that the transition from directed to a mixture of transport and tug of wars occurs well before cargo becomes sterically trapped in the actin network.

### 4 Additional analysis

There are a variety of ways to categorize the motion of liposomes transported by myoVa teams through a given actin network. In the main text, we look at various quantities: the average states of the liposomes in our simulations (Fig. 3) and various quantities that can be measured from liposome trajectories (Fig. 4A-D, average speed, MSD exponent, standard deviation of MSD exponent, and hit targets). Regardless of which quantity we look at, they all show scaling differences on subcritical and supercritical networks, indicating that the percolation phase transition in the actin network causes a change in liposome transport. In this section, we look in more detail at two different ways of characterizing liposome transport. The first, the mode of motion, is a method of differentiating between times that a liposome is stationary, diffusing, or moving in a directed fashion along actin. The second is a method to categorize liposome motion based on measured quantities of liposome transport (average speed, standard deviation of MSD exponent, and hit targets) without any *a priori* assumptions about what types of motion may or may not be present. Regardless of which method we use, our results are the same: a percolation phase transition in the actin network causes a change in liposome transport.

#### 4.1 Modes of motion

The motion of liposomes transported by myoVa through a 3D actin network can be categorized by the probability of observing liposomes in each of three different modes of motion, stationary, diffusive-like, and directed (e.g. Lombardo et al. 2019). These different modes of motion are differentiated based on how mean squared displacement scales with time. A liposome adopts the stationary mode if this scaling exponent, $\alpha$, is between $0 \leq \alpha < 0.66$, the diffusive-like mode if $0.66 \leq \alpha < 1.33$ and the directed mode if $1.33 \leq \alpha \leq 2$. In a dense actin network, a single liposome may adopt each of the three modes of motion at different times, so identifying the modes of motion on a given network requires a method to differentiate the modes (e.g. changepoint analysis).

We simulated the motion of millions of liposomes, all over $\sim 100$ seconds at a sampling rate of 0.05s. It would be a significant computational challenge to analyze all these data with changepoint analysis. However, we can estimate the mode(s) of motion in our simulations by taking advantage of the fact that we have a large amount of data and, because it is simulated data, it is noise-free. To do so, we calculate $\alpha$ for all 5,000 simulated liposomes on a given network over a relatively short time scale (1s) early in the

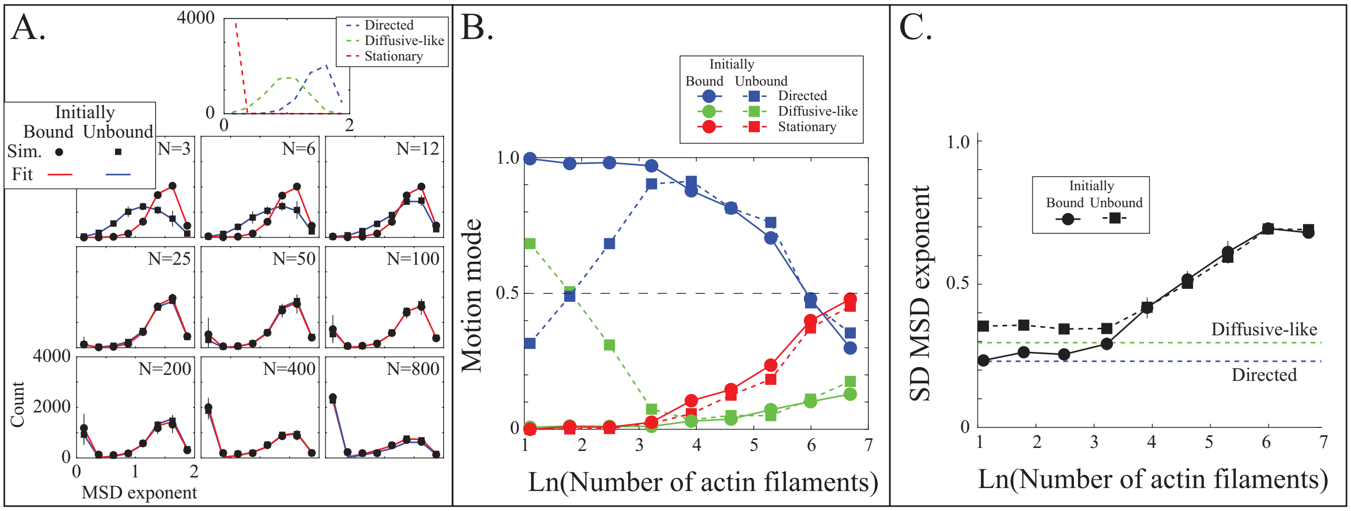


Figure S6: Identifying modes of motion in the simulations. A. Histograms of MSD exponent for different actin network sizes for simulations with liposomes initially unbound (squares) and initially bound (circles). The MSD exponent is calculated from the displacement as a function of time from $t = 9$ to $t = 10$s for all 5,000 simulated liposomes. The dots show averages of simulations on 30 actin networks ($N = 3 - 100$), 15 networks ($N = 200$), 10 networks ($N = 400$) and 5 networks ($N = 800$); error bars show SEM. Lines show best fit combinations of directed, diffusive-like and stationary distributions (top, inset). The directed and diffusive-like distributions were calculated from simulations on actin networks of one and zero filaments, respectively; the stationary distribution is an MSD exponent of 0. B. The proportion of liposomes in each mode of motion as a function of number of actin flaments. Values come from the best-fits shown in A. C. Standard deviation of the distribution of MSD exponent. Squares and dots show averages for simulations with liposomes initially unbound and bound, respectively. Error bars show SEM. This quantity reaches a maximum when the distribution is bimodal, and a minimum when only a single mode of motion occurs.

simulations (between $t = 9$ and $t = 10$s), over which most liposomes will adopt a single mode of motion. The resulting distributions are similar for actin networks of a given size, yet variable over different network sizes or if the liposomes start bound to or unbound from the actin network (Fig. S6A, note the barely distinguishable error bars).

To differentiate the modes of motion, we ran simulations of an actin network with one filament, and with zero filaments. The former gives the expected distribution for the directed mode, while the latter gives the expected distribution for the diffusive-like mode. Note that, over longer time scales, liposomes that are transported along the random actin network would likely have scaling exponents, $\alpha$, in the diffusive-like mode, but we minimize these by picking a relatively short time scale. Finally, stationary liposomes should have $\alpha \approx 0$[3]. We therefore obtain distributions for each of the three modes (Fig. S6A, top).

To determine the proportion of each mode on a given network, we fit the $\alpha$ histogram with a linear combination of the distributions of the three modes, minimizing the $\chi^2$ error. In all cases, the fits are good (Fig. S6A). The coefficients of the linear combination give the relative amounts of each mode (Fig. S6B). In simulations where a myoVa motor starts bound to the actin network, the directed mode dominates at low actin densities (below the percolation phase transition), while the stationary mode becomes more common at higher densities. There is a concurrent rise in diffusive-like motion, likely reflecting the fact that even over this short time scale, liposomes transported along a random actin network can give $\alpha$ values around 1. However, even on the densest actin network, this remains a small proportion ($< 15\%$). In simulations where myoVa motors are unbound from the actin network, the diffusive-like mode dominates at low density,

[3]Note that, unlike in these simulations, in measurements (where small errors in determining liposome location occur) or even in simulations of the detailed model (where the liposome often repositions during a tug of war) liposome position is not exactly constant during a tug of war

but the distributions quickly become nearly indistinguishable from the ones where a liposome starts bound to actin.

We can quantify these transitions by looking at the standard deviation of the $\alpha$ distribution ($\sigma_\alpha$). When the liposomes mostly adopt a single mode of motion, $\sigma_\alpha$ is small; however, when liposomes adopt more than one mode of motion, $\sigma_\alpha$ increases. Therefore, as liposomes switch from directed to a mix of directed and stationary modes (when starting bound to actin), or from diffusive-like to a mix of directed and stationary modes (when starting unbound), we observe and increase in $\sigma_\alpha$ (Fig. S6C). This value reaches a maximum on actin networks of 400 filaments, meaning that the liposomes adopt the stationary and directed modes of motion with roughly equal frequency.

#### 4.2 Categorizing transport

From Fig. 4A of the main text, it is clear that myoVa-driven liposome transport in supercritical actin networks differs from transport on subcritical actin networks. In this section, we quantify this change. To do so, for each simulation of 5,000 liposomes on a given actin network, we looked at three different quantities: 1) average speed, 2) number of targets hit, and 3) the standard deviation of the MSD exponent between $t = 9$ and $t = 10$s (we chose three of the four quantities we measured for visualization purposes). We additionally calculated these quantities for liposomes moving along a single filament, which defines "directed" transport, and for liposomes freely diffusing with no actin, which defines "diffusive" transport.

When we plot these quantities for all of our measurements, we find that many of the simulations cluster near the directed transport and none are near diffusive transport (Fig. S7A). This is consistent with our observation that very little diffusion occurs in our simulations (see Fig. 3 of the main text). However many points are not near directed transport, either. These points represent the random walk trajectories interspersed with pauses that we observe on supercritical networks. To quantify this, we use the simulations on networks of $N = 400$ filaments as a reference since, on these networks, the number of targets hit and the standard deviation of the MSD exponent are both maximized (Fig. 3 of the main text, and Fig. S7A). We refer to this as "mixed" transport, for simplicity.

We can then categorize the transport of 5,000 liposomes on a given actin network by applying a linear scaling to the axes such that each of the three reference points (directed, mixed, diffusive) is at the corner of a unit tetrahedron (i.e., at $(0, 1, 0)$, $(0, 0, 1)$, $(1, 0, 0)$, respectively). Then, we categorize the transport for each simulation as the distance from each corner of the tetrahedron, scaled such that if the simulation is precisely at one corner, it receives a value of 1 for the association transport category (directed, mixed, diffusive) and zeros for the others (Fig. S7A, colors). When plotted against actin network size, we see that there is a transition from directed to mixed that occurs near the percolation threshold, $N_c$ (Fig. S7B).

We applied this same analysis to our simulations with larger, 1,750nm ($r_L = 875$nm) liposomes. The one difference was that, instead of using $N = 400$ for a reference point for mixed transport, we used $N = 100$ since the number of targets hit and the standard deviation of the MSD exponent are maximized on these networks (Fig. 7C of the main text). Like in the simulations with smaller liposomes, these simulations show little diffusion, and motion transitions from directed to mixed transport. Strikingly, when plotted against actin network size and scaled by the percolation threshold, $N_c$, the curves for larger and smaller liposomes are similar (Fig. S7C). This result shows that it is the percolation threshold that determines the change in transport we observe in our simulations.

Finally, we applied this analysis to our simulations with liposomes starting unbound from actin. In these, we observe simulations that cluster near the point that defines diffusive transport (Fig. S7D). The simulations then transition toward the point that defines directed transport before transitioning to mixed transport. Thus, we are able to quantify the changes in myoVa-based liposome transport we observe in our simulations, and the results support our conclusions that 1) as actin density increases, a percolation phase transition drives a change from directed to mixed transport; and 2) this transition occurs whether or not liposomes start with their myoVa bound to actin.

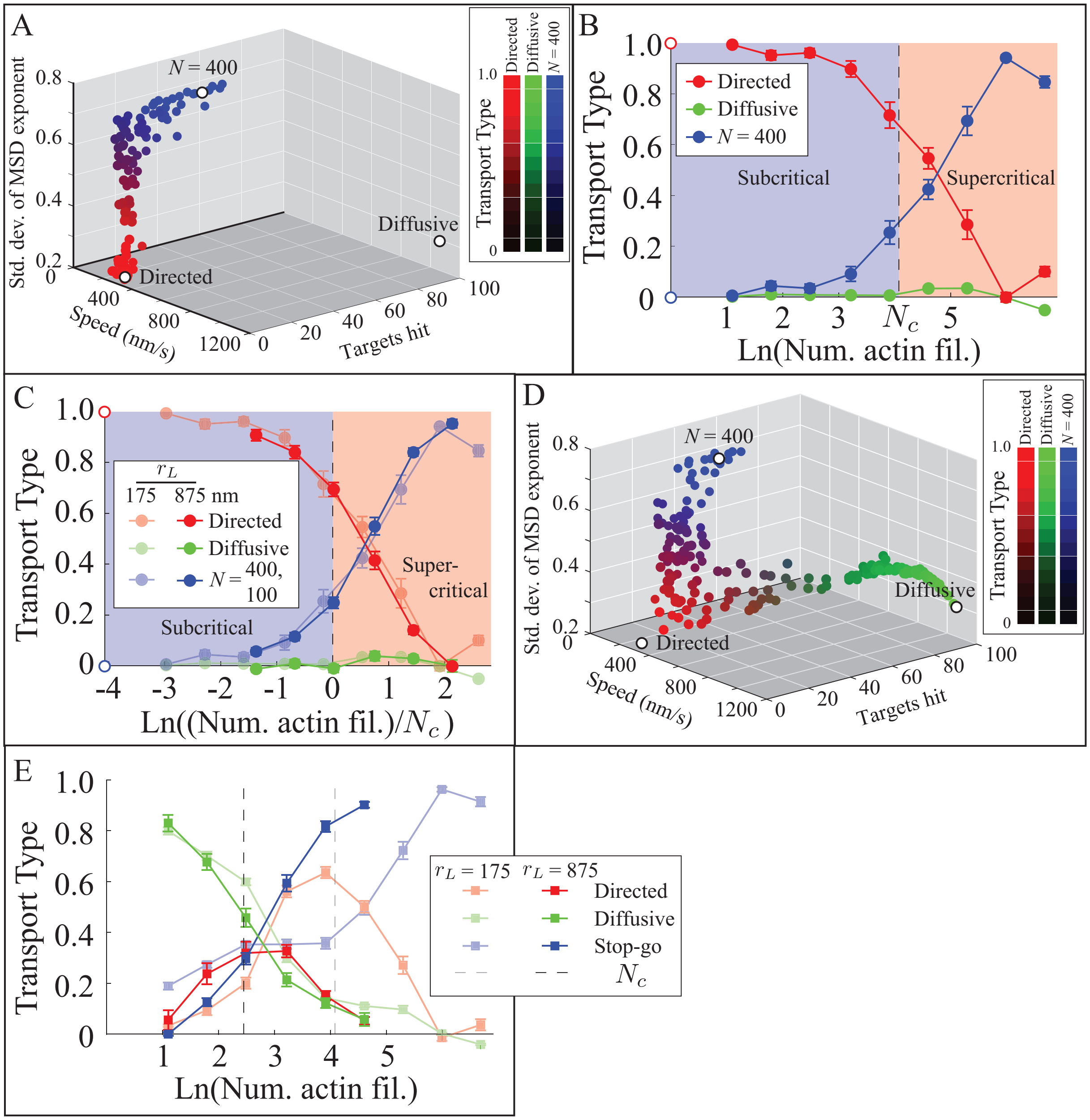


Figure S7: Visualizing the phase transition in transport for small ($r_L = 175$nm) and large ($r_L = 875$nm) liposomes. A. Categorizing transport on a given actin network. Each dot represents the average speed, targets hit and standard deviation of MSD scaling exponent for a single simulation of 5,000 liposomes moving through an actin network. The three hollow circles represent directed transport (observed for liposome transported along a single actin filament), diffusive transport (observed for a liposome in the absence of actin), and $N = 400$ is "mixed" transport. The distance from each of the reference point allows us to quantify transport on a given network (colors). B. Categories of transport as a function of actin density. This is the same data as in A, but shown as a function of the number of actin filaments in the network. C. Categories of transport as a function of actin density for small ($r_L = 175$nm, partially transparent) and large ($r_L = 875$nm, opaque) liposomes are similar when actin density is scaled by the critical actin density, $N_c$. D. Simulations with liposomes initially detached from actin show diffusive transport in addition to the transition from directed to mixed transport. E. The transition from diffusive to directed transport is independent of $N_c$, since it occurs at a similar number of actin filaments for small ($r_L = 175$nm, partially transparent) and large ($r_L = 875$nm, opaque) liposomes. Error bars show SEM.

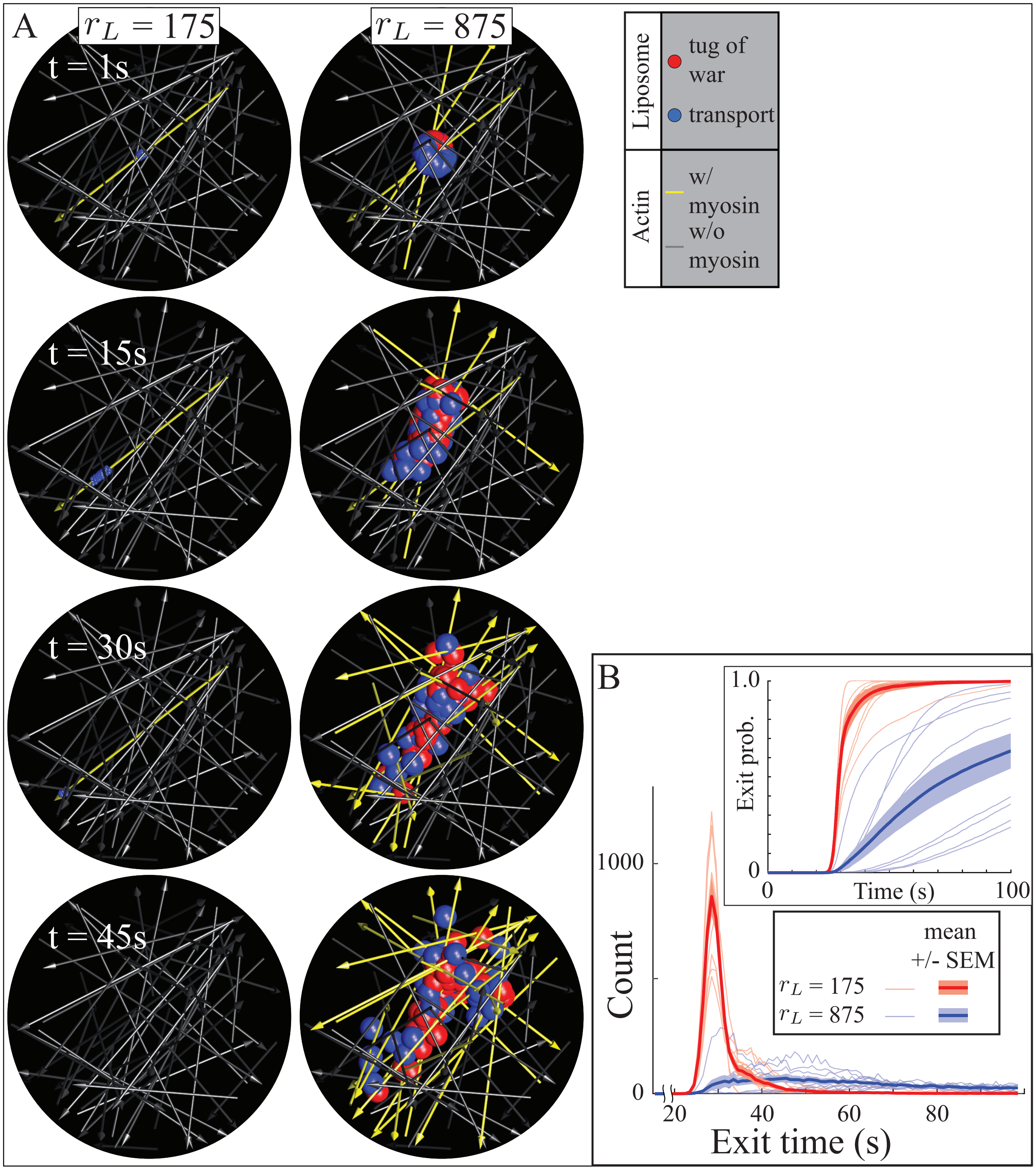


Figure S8: Actin networks can act like a sieve, holding larger liposomes while smaller liposomes are transported. A. Frames from a simulation showing 100 of the 5000 liposomes of $r_L = 175$nm (top) and $r_L = 875$nm (bottom) on an actin network of $N = 50$ filaments. Smaller liposomes are in the transport state (blue) and leave the domain quickly; larger liposomes are in a mix of transport and tug of war (red) states and leave the domain slowly. Actin filament thickness is exaggerated for visualization purposes; arrows indicate plus ends. B. Histograms (bin width 1s) of the number of liposomes leaving the domain show that almost all of the smaller liposomes have left before the larger liposomes begin to leave. Inset shows the probability that a liposome has left the domain as a function of time.

##### 4.3 Actin networks as a sieve

On subcritical networks, we observe directed transport; on supercritical networks, we observe slower trajectories due to liposomes exhibiting a mixture of transport and tug of wars and thus frequently switching actin filaments. Since the phase transition that separates these two types of transport is inversely proportional to liposome radius, on a given actin network sufficiently small liposomes undergo directed transport while sufficiently large liposomes undergo a mix of transport and tug of wars. Such a network acts as a "sieve," though the physical mechanism is different. To demonstrate this behavior, we ran simulations of the smaller ($r_L = 175$nm) and larger ($r_L = 875$nm) liposomes on the same actin networks of 50 filaments, just below the phase transition for the smaller liposomes and well above the phase transition for the larger liposomes. As expected, the smaller liposomes rapidly move to the network boundary sphere to end the simulation, while the larger liposomes move much more slowly (Fig. S8A,B).

#### Supplementary Movies

##### Movie S1

A representative simulation on an actin network of 3 filaments, where the liposomes start with myoVa bound to actin. Left shows the liposomes and the actin network. Right shows the targets and the liposomes. Actin filaments turn yellow once a myoVa motor binds. Liposomes in the transport state are blue, in the tug of war state are red, and in the diffusion state are yellow. Targets are gray until hit by a liposome, then they turn blue. For visualization purposes, only 1,000 of the 5,000 liposomes are shown. Movie is sped up $\sim 6\times$ real time.

##### Movie S2

A representative simulation on an actin network of 25 filaments. Left shows the liposomes and the actin network. Right shows the targets and the liposomes. Actin filaments turn yellow once a myoVa motor binds. Liposomes in the transport state are blue, in the tug of war state are red, and in the diffusion state are yellow. Targets are gray until hit by a liposome, then they turn blue. For visualization purposes, only 1,000 of the 5,000 liposomes are shown. Movie is sped up $\sim 6\times$ real time.

##### Movie S3

A second representative simulation on an actin network of 25 filaments, where the liposomes start with myoVa bound to actin. Left shows the liposomes and the actin network. Right shows the targets and the liposomes. Actin filaments turn yellow once a myoVa motor binds. Liposomes in the transport state are blue, in the tug of war state are red, and in the diffusion state are yellow. Targets are gray until hit by a liposome, then they turn blue. For visualization purposes, only 1,000 of the 5,000 liposomes are shown. Movie is sped up $\sim 6\times$ real time.

#### Movie S4

A representative simulation on an actin network of 50 filaments, where the liposomes start with myoVa bound to actin. Left shows the liposomes and the actin network. Right shows the targets and the liposomes. Actin filaments turn yellow once a myoVa motor binds. Liposomes in the transport state are blue, in the tug of war state are red, and in the diffusion state are yellow. Targets are gray until hit by a liposome, then they turn blue. For visualization purposes, only 1,000 of the 5,000 liposomes are shown. Movie is sped up $\sim 6\times$ real time.

#### Movie S5

A representative simulation on an actin network of 100 filaments, where the liposomes start with myoVa bound to actin. Left shows the liposomes and the actin network. Right shows the targets and the liposomes. Actin filaments turn yellow once a myoVa motor binds. Liposomes in the transport state are blue, in the tug of war state are red, and in the diffusion state are yellow. Targets are gray until hit by a liposome, then they turn blue. For visualization purposes, only 1,000 of the 5,000 liposomes are shown. Movie is sped up $\sim 6\times$ real time.

#### Movie S6

A representative simulation on an actin network of 200 filaments, where the liposomes start with myoVa bound to actin. Left shows the liposomes and the actin network. Right shows the targets and the liposomes. Actin filaments turn yellow once a myoVa motor binds. Liposomes in the transport state are blue, in the tug of war state are red, and in the diffusion state are yellow. Targets are gray until hit by a liposome, then they turn blue. For visualization purposes, only 1,000 of the 5,000 liposomes are shown. Movie is sped up $\sim 6\times$ real time.

#### Movie S7

A simulation on an actin network of 25 filaments, relating to the images shown in Fig. 1. Note that 100 liposomes are shown in the figure, and 1,000 are shown here. Left shows the liposomes and the actin network. Right shows the targets and the liposomes. Actin filaments turn yellow once a myoVa motor binds. Liposomes in the transport state are blue, in the tug of war state are red, and in the diffusion state are yellow. Targets are gray until hit by a liposome, then they turn blue. Movie is sped up $\sim 6\times$ real time.

#### Movie S8

A simulation on an actin network of 50 filaments, relating to the plots shown in Fig. 4F and S4. In this movie, unlike the previous movies, actin filaments are yellow only as long as a myoVa motor is bound, and revert to gray once no myoVa are bound. Left shows the liposomes and the actin network. Right shows the targets and the liposomes. Liposomes in the transport state are blue, in the tug of war state are red, and in the diffusion state are yellow. Targets are gray until hit by a liposome, then they turn blue. Movie is sped up $\sim 12\times$ real time.

#### Movie S9

A simulation on an actin network of 100 filaments, relating to the plots shown in Fig. 4F. In this movie, like Movie S8, actin filaments are yellow only as long as a myoVa motor is bound, and revert to gray once no myoVa are bound. Left shows the liposomes and the actin network. Right shows the targets and the liposomes. Liposomes in the transport state are blue, in the tug of war state are red, and in the diffusion state are yellow. Targets are gray until hit by a liposome, then they turn blue. Movie is sped up $\sim 12\times$ real time.

##### Movie S10

A representative simulation on an actin network of 3 filaments, where the liposomes start with myoVa unbound from actin. Left shows the liposomes and the actin network. Right shows the targets and the liposomes. Actin filaments turn yellow once a myoVa motor binds. Liposomes in the transport state are blue, in the tug of war state are red, and in the diffusion state are yellow. Targets are gray until hit by a liposome, then they turn blue. For visualization purposes, only 1,000 of the 5,000 liposomes are shown. Movie is sped up $\sim 6\times$ real time.

##### Movie S11

A representative simulation on an actin network of 25 filaments, where the liposomes start with myoVa unbound from actin. Left shows the liposomes and the actin network. Right shows the targets and the liposomes. Actin filaments turn yellow once a myoVa motor binds. Liposomes in the transport state are blue, in the tug of war state are red, and in the diffusion state are yellow. Targets are gray until hit by a liposome, then they turn blue. For visualization purposes, only 1,000 of the 5,000 liposomes are shown. Movie is sped up $\sim 6\times$ real time.

##### Movie S12

A representative simulation on an actin network of 50 filaments, where the liposomes start with myoVa unbound from actin. Left shows the liposomes and the actin network. Right shows the targets and the liposomes. Actin filaments turn yellow once a myoVa motor binds. Liposomes in the transport state are blue, in the tug of war state are red, and in the diffusion state are yellow. Targets are gray until hit by a liposome, then they turn blue. For visualization purposes, only 1,000 of the 5,000 liposomes are shown. Movie is sped up $\sim 6\times$ real time.

##### Movie S13

A representative simulation on an actin network of 100 filaments, where the liposomes start with myoVa unbound from actin. Left shows the liposomes and the actin network. Right shows the targets and the liposomes. Actin filaments turn yellow once a myoVa motor binds. Liposomes in the transport state are blue, in the tug of war state are red, and in the diffusion state are yellow. Targets are gray until hit by a liposome, then they turn blue. For visualization purposes, only 1,000 of the 5,000 liposomes are shown. Movie is sped up $\sim 6\times$ real time.

##### Movie S14

A representative simulation on an actin network of 200 filaments, where the liposomes start with myoVa unbound from actin. Left shows the liposomes and the actin network. Right shows the targets and the liposomes. Actin filaments turn yellow once a myoVa motor binds. Liposomes in the transport state are blue, in the tug of war state are red, and in the diffusion state are yellow. Targets are gray until hit by a liposome, then they turn blue. For visualization purposes, only 1,000 of the 5,000 liposomes are shown. Movie is sped up $\sim 6\times$ real time.

#### Movie S15

One simulation shown in Fig. S1B of 50 liposomes encountering an actin "intersection." A top view of two actin filaments (yellow and red lines), separated by 100nm. Horizontal and vertical axes are labeled in nm. The center of the liposomes are pictured as: red circles (no outcome assigned), black dots (straight outcome), blue dots (turn outcome), red dots (terminate outcome). The interaction zone is shown as a pink rectangle. Movie plays approximately real time. Note that this simulation keeps track of steric interactions between the liposome and actin, the myoVa motors and their interactions with actin, and the location of binding sites on actin, although they are not pictured here (see [1] for details and movies that show all elements of the model).

#### Movie S16

A visualization of the pore calculation shown in Fig. S5A of a network of 100 actin filaments. The movie first shows the set of spheres that define the pore in blue. Then, the smallest of those spheres is shown in white, and the camera follows it as it navigates through the pore. The path through the sphere is shown with a gray striped line. The movie then zooms out and shows the final position of the sphere and the path it followed. Actin filament thickness is exaggerated for visualization purposes.
